# Gestational CBD shapes insular cortex in adulthood

**DOI:** 10.1101/2024.06.26.597499

**Authors:** Daniela Iezzi, Alba Cáceres-Rodríguez, Jessica Pereira Silva, Pascale Chavis, Olivier J. Manzoni

## Abstract

Many expectant mothers use CBD to alleviate symptoms like nausea, insomnia, anxiety, and pain, despite limited research on its long-term effects. However, CBD passes through the placenta, affecting fetal development and impacting offspring behavior. We investigated how prenatal CBD exposure affects the insular cortex (IC), a brain region involved in emotional processing and linked to psychiatric disorders. The IC is divided into two territories: the anterior IC (aIC), processing socioemotional signals, and the posterior IC (pIC), specializing in interoception and pain perception. Pyramidal neurons in the aIC and pIC exhibit sex-specific electrophysiological properties, including variations in excitability and the excitatory/inhibitory balance. We investigated IC’s cellular properties and synaptic strength in the offspring of both sexes from mice exposed to low-dose CBD during gestation (E5-E18; 3mg/kg, s.c.). Prenatal CBD exposure induced sex-specific and territory-specific changes in the active and passive membrane properties, as well as intrinsic excitability and the excitatory/inhibitory balance, in the IC of adult offspring. The data indicate that in-utero CBD exposure disrupts IC neuronal development, leading to a loss of functional distinction between IC territories. These findings may have significant implications for understanding the effects of CBD on emotional behaviors in offspring.

## 1. Introduction

The growing trend of cannabis use among expectant and nursing mothers is sparking mounting alarm, posing a substantial threat to the well-being of unborn babies and newborns. The two main psychoactive compounds in cannabis, tetrahydrocannabinol (THC) and cannabidiol (CBD), can permeate the placenta during pregnancy (as extensively discussed in Rokeby et al. [1]) and also concentrate in breast milk during lactation. This raises concerns about the potential risks of in utero exposure to these substances and their impact on the vulnerable developing fetus and breastfeeding infant. While CBD shares a structural similarity to Δ9-THC, it is often advertised as a non-psychotropic substance. Consequently, CBD from processed hemp products such as CBD oils and extracts is generally perceived as safe and free of adverse effects.

Although there is a dearth of scientific research on the safety of CBD during pregnancy, expectant mothers are increasingly turning to CBD to alleviate a range of pregnancy-related complaints, including morning sickness, sleep disturbances, anxiety, and persistent pain [2]. Moreover, CBD has been shown to penetrate the human placenta, which may have implications for the developing fetus and the functioning of the placenta itself [1], [3]–[5]. Additionally, CBD (and THC) concentrations are elevated in the plasma and breast milk of lactating mothers [6], which may have implications for the nursing infant. Of particular significance, CBD [7] can exert a profound impact on the differentiation, maturation, and functionality of human induced pluripotent stem cells, suggesting a potential neurodevelopmental effect on the developing fetus and infant.

In rodent models too, CBD has also been shown to traverse the placenta and accumulate in breast milk [8]. We previously showed that administration of a low dose of CBD (3mg/kg) from embryonic day 5 to 18 (E5-E18) during gestation resulted in sex-specific behavioral alterations in neonatal pups. Specifically, male pups exhibited increased weight gain and emitted shorter ultrasonic vocalizations when separated from the nest, whereas female littermates produced more high-frequency calls. Notably, the qualitative changes observed in the syllabic repertoire of ultrasonic vocalizations suggest altered communication patterns in the offspring exposed to prenatal CBD. Furthermore, female pups exposed to CBD during gestation displayed reduced motor and discriminatory abilities, indicating a higher susceptibility to the effects of prenatal CBD compared to males, which may have implications for the long-term development and behavior of the offspring [9].

The repercussions of perinatal CBD exposure persist beyond the early life stages. Continuous exposure to CBD from gestation until the first week postpartum led to sex-specific alterations in repetitive and hedonic behaviors in adult offspring [10]. Moreover, prenatal CBD exposure was found to attenuate the ability of fluoxetine to enhance coping behavior in the forced-swim test, a paradigm used to assess antidepressant-like effects [10].

The insular cortex (IC) plays a pivotal role in various cognitive and emotional processes. It is involved in sensory processing, representing feelings and emotions, autonomic and motor control, risk prediction, decision-making, bodily and self-awareness, as well as complex social functions such as empathy [11], [12]. The IC functions as an integration hub by connecting diverse brain systems underlying sensory, emotional, motivational, and cognitive functions[13]. Thus, the IC receives sensory inputs from various modalities, projects to topographically organized sensory regions, and connects with the limbic system, frontal brain regions, and regions implicated in motivation and reward [14], [15]. The IC is also associated with interoception, emotional valence, decision-making under uncertainty, empathy, and even self-awareness. The insular IC has been associated with a range of neurological and neuropsychiatric disorders: alterations or dysfunctions in the IC have been linked to anxiety disorders, addiction, depression, schizophrenia, and autism spectrum disorders [16].

The IC s divided into two territories. The anterior insular cortex (aIC) plays a critical role in the processing and experience of emotions, social interactions, and self-awareness [12], while the posterior insular cortex pIC is associated with sensory integration, interoceptive awareness, and motor control [13], [17]. In a preceding investigation, we characterized and compared the cellular and synaptic properties of pyramidal neurons across IC territories in mice of both sexes, and revealed region-specific electrophysiological signatures and sex-dependent differences in synaptic plasticity and excitatory transmission [18]. These regional disparities likely contribute to the distinct behavioral and cognitive profiles observed between males and females, and may underlie the specialized functions of the aIC and pIC.

Here, we examined the cellular characteristics and synaptic strength of the IC in both male and female progeny of mice subjected to a low dose of CBD (3mg/kg, s.c.) during gestation (E5-E18). The results show that prenatal CBD exposure triggers sex-specific and territory-specific modifications in the electrophysiological properties, as well as intrinsic excitability and the balance of excitatory and inhibitory inputs, in the IC of adult progeny. In-utero CBD leads to a diminution of the normal functional heterogeneity between the IC’s territories. These findings have important implications for our understanding of the effects of CBD on behaviors in offspring.

## 2. Materials and methods

### 2.1 Animals

Animals were treated in compliance with the European Communities Council Directive (86/609/EEC) and the United States NIH Guide for the Care and Use of Laboratory Animals. The French Ethical committee authorized the project APAFIS#18476-2019022510121076 v3. Adult male and female C57BL6/J (12–17 weeks age) were purchased from Charles River and housed in standard wire-topped Plexiglas cages (42□×□27□x□14□cm), in a temperature and humidity-controlled condition (i.e., temperature 21□±□1□°C, 60□±□10% relative humidity and 12□h light/dark cycles). Food and water were available ad *libitum*. Following a one-week acclimation period, female pairs were introduced to a single male mouse in the late afternoon. The day when a vaginal plug was observed was considered as day 0 of gestation (GD0), and pregnant mice were individually housed thereafter. Starting from GD5 until GD18, the pregnant dams received daily subcutaneous injections (s.c.) of either a vehicle or 3 mg/kg of CBD (obtained from the Nida Drug Supply Program). The CBD was dissolved in a vehicle solution composed of Cremophor EL (from Sigma-Aldrich), ethanol, and saline, with ratios of 1:1:18, respectively, and administered at a volume of 4 mL/kg. Control dams (referred to as "Sham") received an equivalent volume of the vehicle solution. Upon each litter’s birth, the day was designated as postnatal day (PND) 0. Pups were weaned on postnatal day (PND) 21 and subsequently housed separately by sex.

### 2.2 Slice preparation

Adult male and female mice (PND 90-120) were deeply anesthetized with isoflurane and sacrificed according to institutional regulations. The brain was sliced (300□μm) in the coronal plane with a vibratome (Integraslice, Campden Instruments) in a sucrose-based solution at 4°C (87 mM NaCl, 75 mM sucrose, 25 mM glucose, 2.5 mM KCl, 4 mM MgCl2, 0.5 mM CaCl2, 23 mM NaHCO3, and 1.25 mM NaH2PO4). Immediately after cutting, slices containing anterior or posterior IC were stored for 30 minutes at 32°C in a low-calcium artificial CSF (ACSF) that contained the following: 130 mM NaCl, 11 mM glucose, 2.5 mM KCl, 2.4 mM MgCl2, 1.2 mM CaCl2, 23 mM NaHCO3, and 1.2 mM NaH2PO4, and were equilibrated with 95% O2/5% CO2 and then at room temperature until the time of recording. During the recording, slices were placed in the recording chamber and continuously perfused at 2 ml/min with warm (32°-34°C) low Ca2+ solution.

### 2.3 Electrophysiology

Whole-cell patch clamp recordings were made from the soma of layer V pyramidal anterior or posterior IC neurons. The latter were visualized under a differential interference contrast microscope using an upright microscope with infrared illumination (Olympus, France). For current-clamp experiments and voltage clamp recording, patch pipettes were filled with an intracellular solution containing (in mM): K^+^ gluconate (145 K^+^ gluconate, 3 NaCl, 1 MgCl_2_, 1 EGTA, 0.3 CaCl_2_, 2 Na^2+^ ATP, 0.3 Na^+^ GTP, and 0.2 cAMP, buffered with 10 HEPES). The pH was adjusted to 7.25 and osmolarity to 290 – 300 mOsm. Electrode resistance was 2 – 4 MΩ. Access resistance compensation was not used, and acceptable access resistance was <30 MΩ. The potential reference of the amplifier was adjusted to zero before breaking into the cell. Cells were held at –70 mV. Current-voltage (I-V) curves were made by a series of hyperpolarizing to depolarizing current steps immediately after breaking into the cell. To determine rheobase a series of depolarizing current steps was applied. Spontaneous EPSCs (sEPSCs) were recorded at −70 mV and isolated by using the GABA_A_ receptor blocker gabazine 10 mM (SR 95531 hydrobromide; Tocris).

When inhibitory postsynaptic currents (IPSCs) were recorded, the recording pipettes were filled with a high-chloride solution of the following composition (in mM): 140 KCl, 1.6 MgCl_2_, 2.5 MgATP, 0.5 NaGTP, 2 EGTA, 10 HEPES. The pH solution was adjusted to 7.25-7.3 and osmolarity to 280 – 300 mOsm. Electrode resistance was 3–4 MΩ. Spontaneous IPSCs (sIPSCs) were recorded at −70mV in presence of 10□μM CNQX (6-Cyano-7-nitroquinoxaline-2,3-dione disodium, an AMPA receptor antagonist, Tocris) and L-APV 50 μM (DL-2-Amino-5-phosphonopentanoic acid, a selective NMDA receptor antagonist, Tocris).

Data was recorded in current clamp with an Axopatch-200B amplifier, low pass filtered at 2 kHz, digitized (10 kHz, DigiData 1440A, Axon Instruments), collected and analyzed using Clampex 10.7 (Molecular Device).

### 2.4 Data analysis and statistics

Except for Principal Component Analysis (PCA, see below), data were analysed off-line with Clampfit 10.7 (Molecular Devices, Sunnyvale, CA, USA) and AxoGraph X. Graphs and Figure layouts were generated with GraphPad Prism 10. Datasets were tested for the normality (D’Agostino-Pearson and Shapiro-Wilk) and outliers (ROUT test) before running parametric tests. Statistical significance of difference between means was assessed with two-way ANOVA followed by Sidak’s multiple comparison post hoc tests as indicated in figure legends. When achieved, the significance was expressed as exactly *p*-value in the figures. The experimental results are described qualitatively in the main text, whereas experimental and statistical details, including sample size (n/N = Cells/Animals), statistical test, *p*-value, main effects and interactions, are reported in Figure legends or in Supplementary Tables. Quantitative data are presented as Box and whisker plots, reporting median, min and max values, and superimposed scatter plots to show individual data points. Membrane capacitance (Cm) was estimated by integrating the capacitive current evoked by a −2 mV pulse whereas the membrane resistance was estimated from the I-V curve around resting membrane potential. The latter (RMP) was measured immediately after whole-cell formation during the current-clamp protocol. The input-output curve was made by measuring the number of action potentials elicited by depolarizing current steps of increasing amplitude, while to determine rheobase a series of depolarizing 10 pA current steps was applied. The frequency and amplitude of sE/IPSCs were analysed with Axograph X using a double exponential template: f(t) = exp(-t/rise) + exp(-t/decay) (rise = 0.5 ms and decay = 3 ms; rise = 0.2 ms and decay = 10 ms, respectively). The detection threshold for the events was set at 3 times the baseline noise SD, whereas the one for the amplitude detection was set at -7 pA. The radar plot was constructed using the electrophysiological properties of principal neurons in aIC and pIC, considering both sex and treatment variables. The means of each population were normalized within a range of -2 to 5 using the following function: Normalized Values = -2 + (Values - Minimum) * (-5 - (-2))/(Maximum - Minimum).

Correlations were compared with “cocor.indep.groups” of the package cocor [19] in R 4.2.0 [20]. Comparison results were shown only for those correlations found to be significant in at least one of the two compared groups.

Considering the parameters reported in the evaluation of intrinsic and synaptic transmission properties, Principal Component Analysis (PCA) was computed using “The FactoMineR package”. Intrinsic properties of layer 5 pyramidal neurons were analysed via PCA with membrane capacitance, rheobase, resting membrane potentials, neuronal excitabilities and voltage membrane response to a different injected current steps as quantitative variables and individual cells as individuals. Missing values were imputed using the mean of the respective variable. Supplementary qualitative variables were the two insular cortices (anterior and posterior, 2 modalities), treatment (Sham and CBD, 2 modalities) and group (4 modalities). The cumulative relative contribution of PCs against the variance, the contribution and correlation were investigated.

## 3. Results

### 3.1 Gestational CBD induced sex- and territory-specific alterations in the properties of IC pyramidal neurons

Prenatal environmental risk factors can profoundly disrupt brain development and function in the offspring, thereby increasing susceptibility to a range of neurodevelopmental and neuropsychiatric disorder [21]. The IC integrates multiple sensory modalities and the processes emotional stimuli. Considering its significance in the development of psychiatric conditions, we examined the impact of prenatal CBD exposure on the functionality of the IC. We previously observed significant differences in the cellular properties of layer V pyramidal output neurons along the rostro-caudal axis of the IC in adult males and females [18]. We characterized the passive and active membrane properties of layer V pyramidal neurons in the aIC and pIC of CBD-exposed progeny, examining both sexes (Figures 2 and 3; Tables 1 and 2, Supplementary Figures 1 and 2). In males exposed to CBD, there were no differences in soma size and resting membrane potential between the aIC and pIC regions, in contrast with the findings in the Sham group and Naïve mice [18] (Figure 2A-B). However, females exposed to CBD exhibited larger and more hyperpolarized pyramidal neurons like Sham females across IC subregions (Figure 2D-E). Furthermore, both sexes of CBD-exposed mice showed a comparable rheobase between the aIC and pIC (Figure 2C, F). When comparing passive and active membrane properties within each IC, we observed that pIC pyramidal neurons were more hyperpolarized in CBD-exposed males compared to their Sham counterparts (Figure 2B). Additionally, the rheobase of pIC neurons was higher in both CBD-exposed males and females compared to the Sham groups, indicating a more negative resting membrane potential (Figure 2C, F).

**Figure 1.**
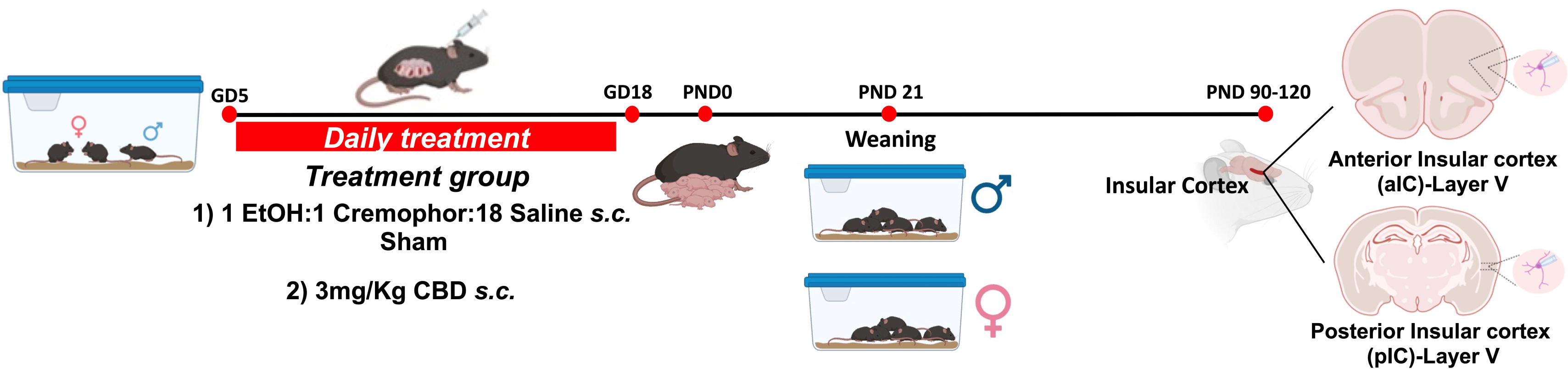
Experimental timeline for gestational *Cannabidiol* (CBD) exposure. Mice were bred and treated once daily from gestational day (GD) 5 to GD18. The date of born was defined as post-natal day (PND) 0. Pups were weaned at PND 21 and male and female were housed separately. In adult stage electrophysiological recordings were made on layer V principal neurons of anterior and posterior Insular Cortex (aIC and pIC respectively), in both sexes of Sham and CBD exposed progeny. Created with Biorender.com.

**Figure 2.**
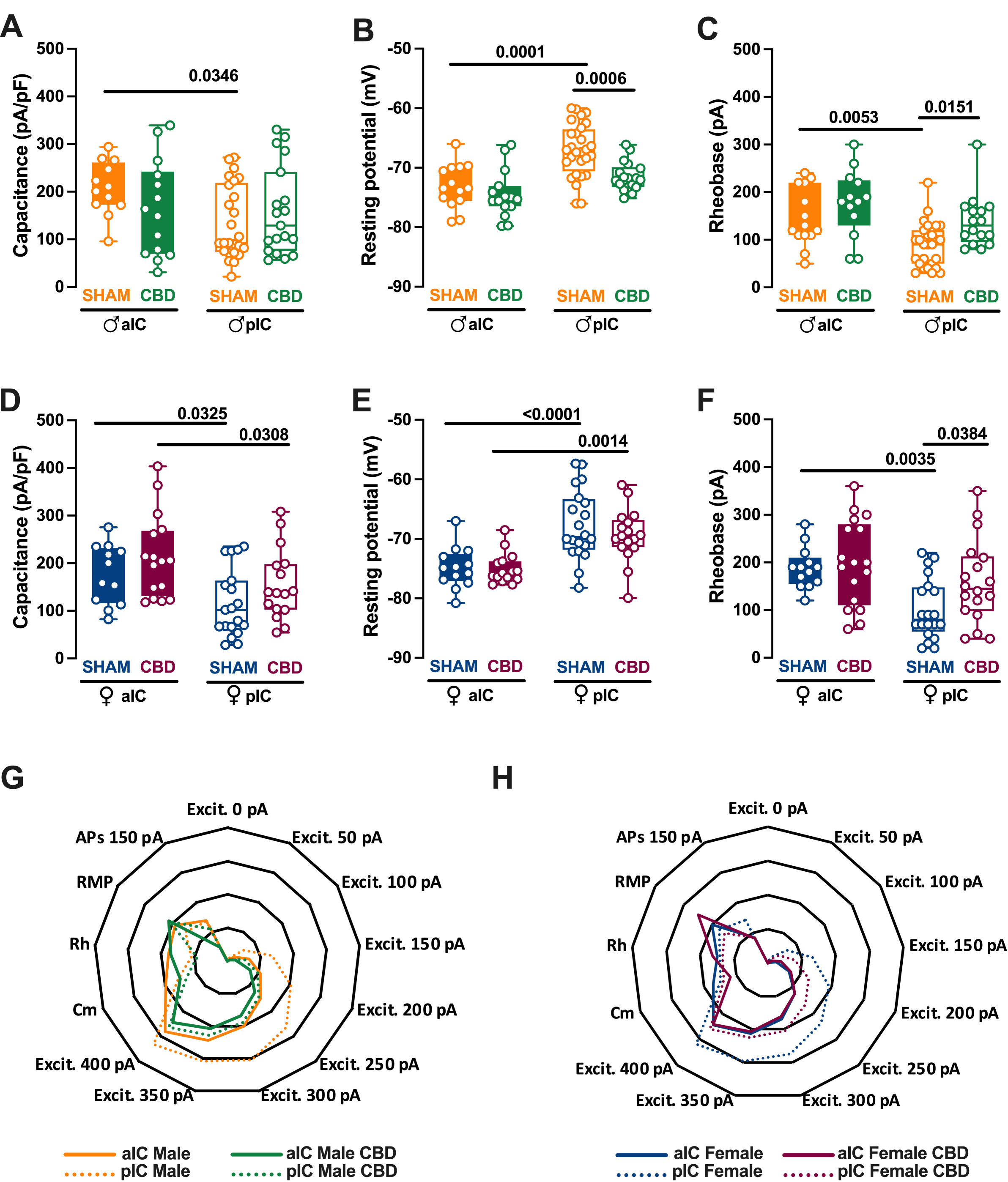
Gestation CDB exposure induces a subregion specific alteration of intrinsic properties of layer V IC pyramidal neurons. **(A-B, D-E)** Quantitative analysis of passive and active membrane properties revealed that across Insular cortex (IC) subregions, principal neurons of anterior IC (aIC) are significantly larger (i.e., larger capacitance) and hyperpolarized compared to posterior IC (pIC) neurons in Sham mice in both sexes. In contrast, following CBD prenatal exposure this difference was completely lost in CBD male, but not in CBD female. Resting membrane potential of pIC neurons is more hyperpolarized in only CBD male compared to the Sham counterpart. **(C, F)** Sham mice showed a higher rheobase in aIC compared to pIC pyramidal neurons in both sexes, but in CBD exposed mice the rheobase was similar across IC subregions while higher in only pIC of both male and female. **(G-H)** Radar plot shows differences between aIC and pIC pyramidal neurons across all electrophysiological properties extracted from Sham and CBD mice, in both sexes. Sham mice showed a clear differentiation in the cellular properties of aIC and pIC, in both male and female. In contrast, following CBD in-utero exposure, across IC subregions, pIC totally lost its specificity, in both sexes. Data are presented as box-and-whisker plots (minimum, maximum, median) for **(A-F)**, and as normalized in a range of -2 and 5 for **(G-H)**. Two-way ANOVA followed by Šídák’s multiple comparison test was performed for **(A-F)**. P– values < 0.05 depicted in the graph. aIC Sham male = 14/10, pIC Sham male = 27/15, aIC Sham female = 14/6, pIC Sham female = 19/13, aIC CBD male = 15/8, pIC CBD male = 19/9, aIC CBD female = 17/7, pIC CBD female = 17/6.

**Figure 3:**
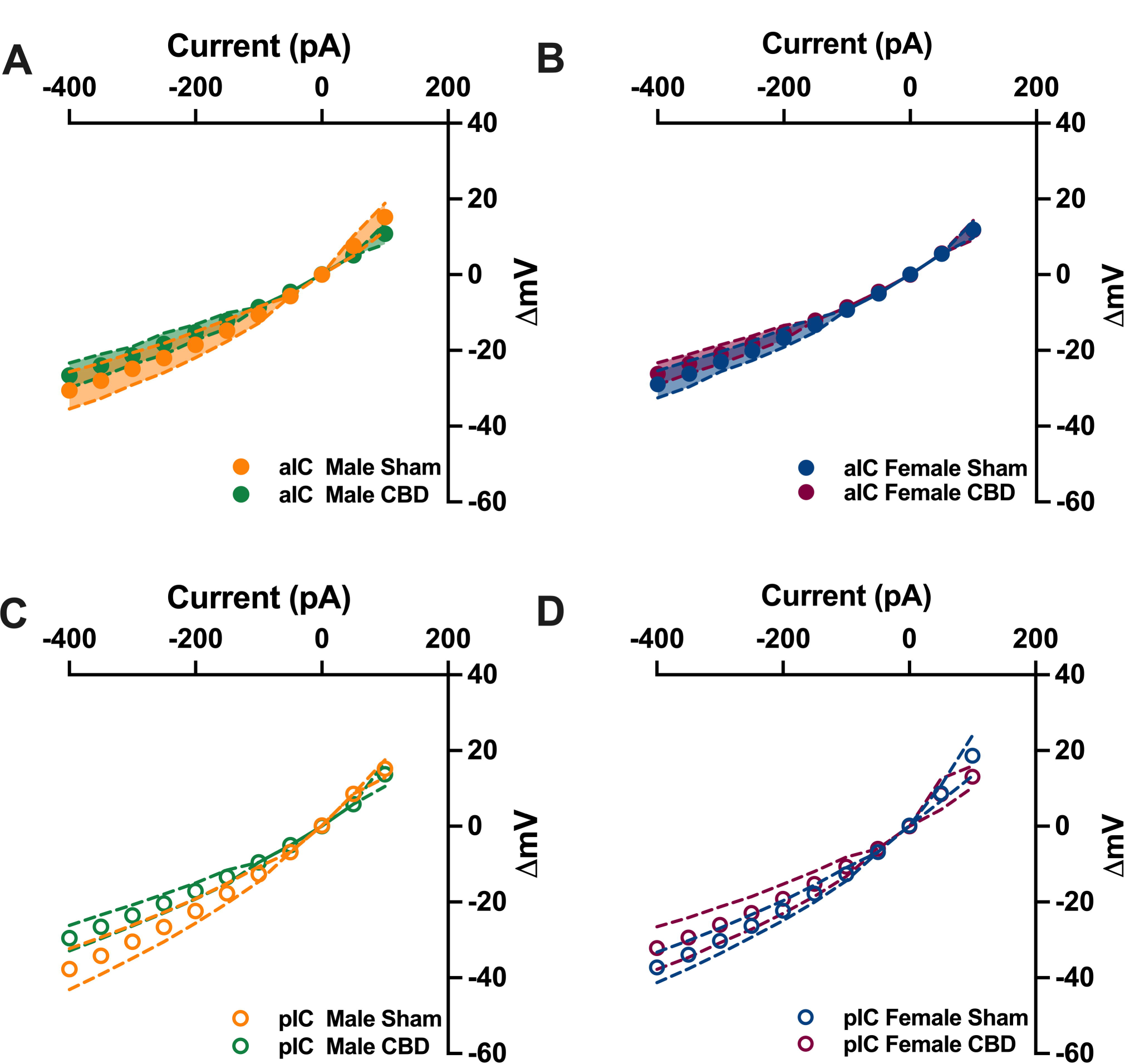
Comparative analysis of I-V relationships across IC subregions following in utero CBD exposure. Panels **A-D** show the I-V curves for each subregion, plotted as mean ± CI. The analysis reveals that CBD exposure in utero does not significantly alter the I-V relationships in IC subregions, regardless of sex. aIC Sham male = 14/10, pIC Sham male = 27/15, aIC Sham female = 14/6, pIC Sham female = 19/13, aIC CBD male = 15/8, pIC CBD male = 19/9, aIC CBD female = 17/7, pIC CBD female = 17/6.ence interval (XY plots). Statistical analysis was performed using the Mann-Whitney U test, with *p < 0.05 considered significant.

**Table 1.**
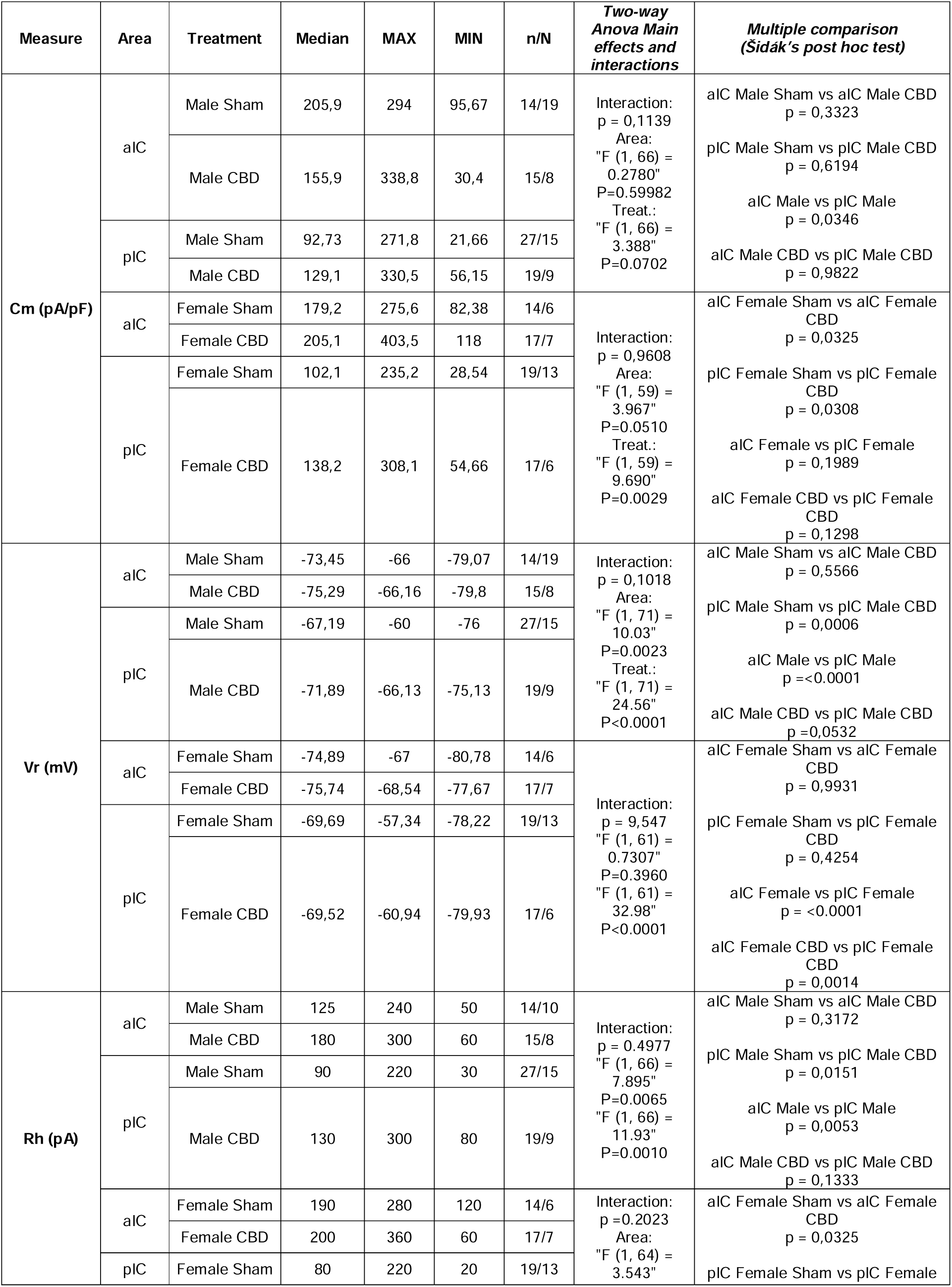
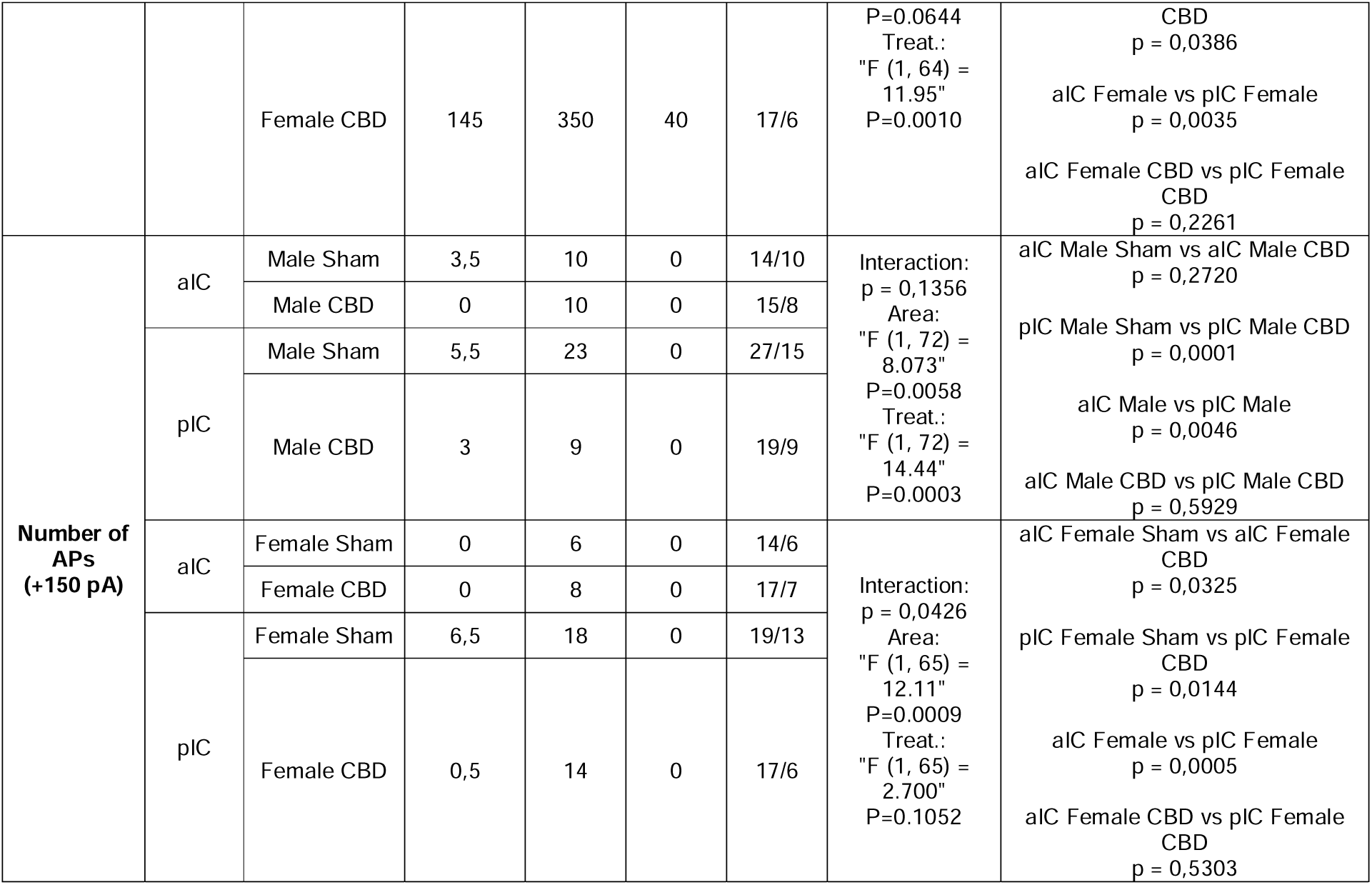
Active and passive membrane properties of aIC and pIC pyramidal neurons across IC subregions and sex. Significance defined as p-value < 0.05.

**Table 2.**
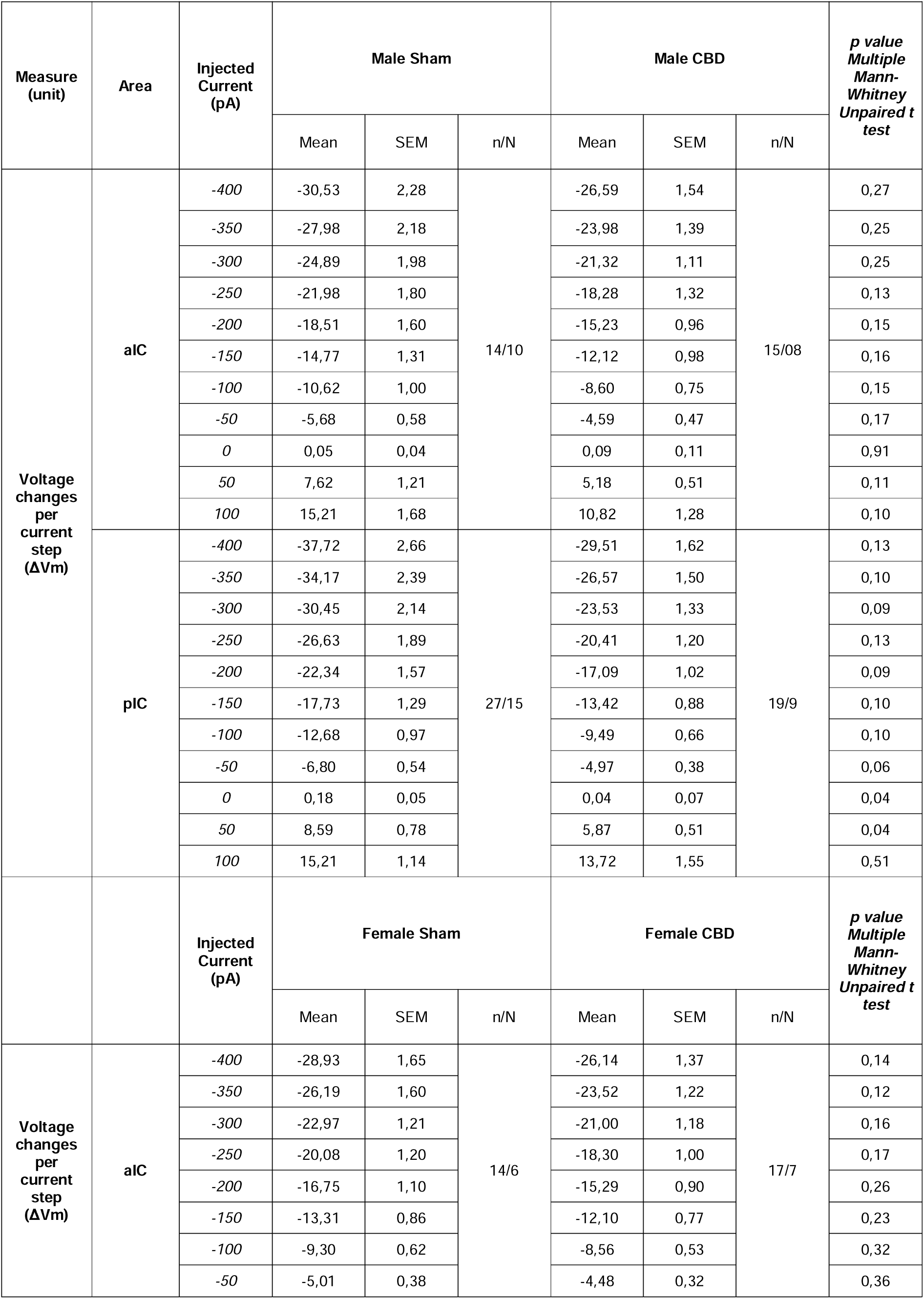
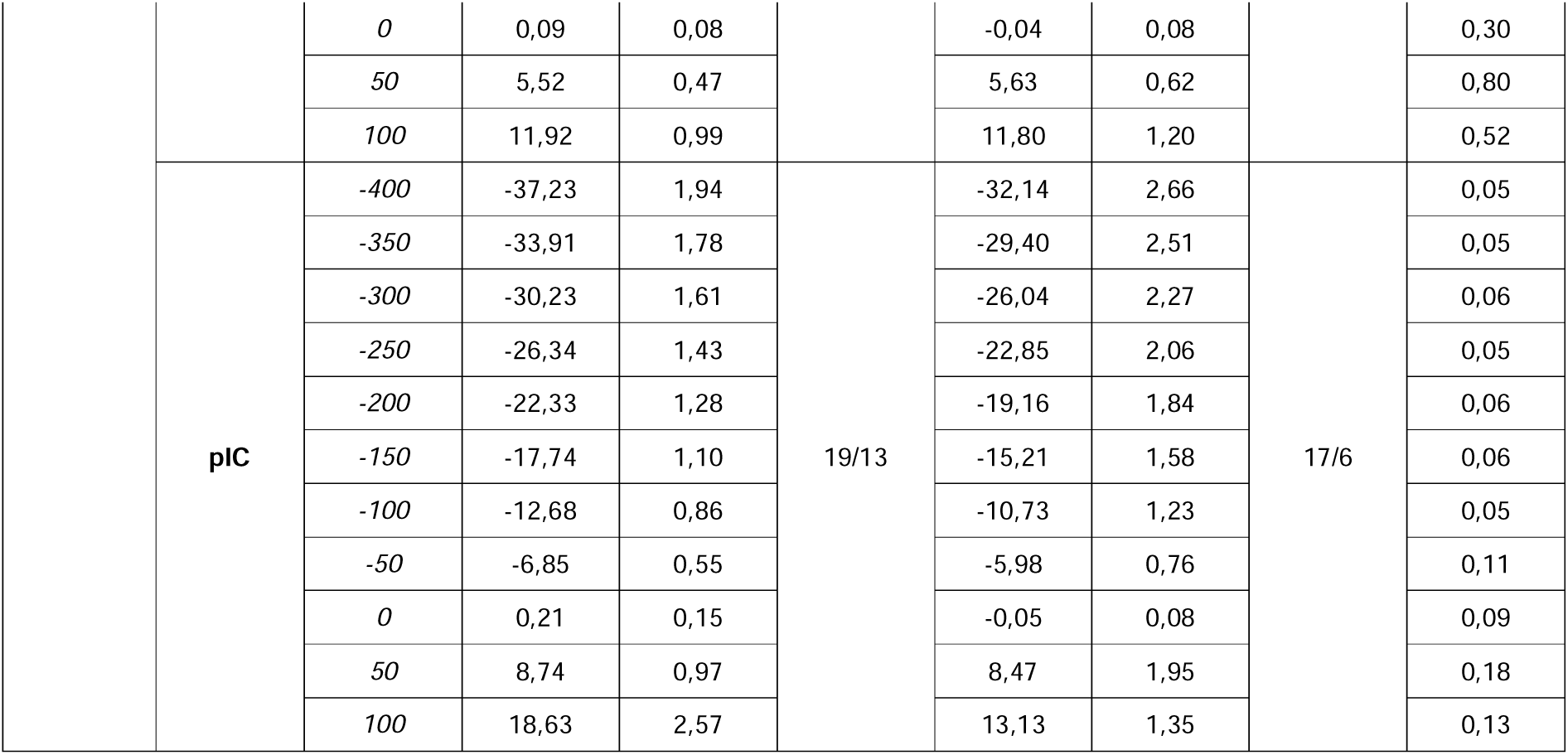
Voltage membrane response per current steps in aIC and pIC pyramidal neurons in both sexes and
treatment. Significance defined as p-value < 0.05.

The comparison of voltage membrane response to hyperpolarizing current steps revealed no significant differences within IC subregions across treatments (Figure 3A-D). Notably, in females (but not in males), pyramidal neurons in the aIC were larger and more hyperpolarized compared to those in the pIC (Supplementary Figure 1A-B). Additionally, irrespective of sex and IC subregion, CBD-exposed progeny displayed similar rheobase values and membrane responses to somatic injection current steps (Supplementary Figure 1C and 2A-D).

Summarizing the cellular properties of principal neurons in the IC across sex and subregions (Figure 2G-H), we observed that fetal CBD exposure specifically disrupted the differentiation of the pIC in CBD-exposed progeny of both sexes. In CBD mice, the active and membrane properties between the aIC and pIC largely overlapped, whereas in the Sham group, these properties exhibited significant differences along the antero-posterior axes for both males and females. Furthermore, we conducted Principal Component Analysis (PCA, Figure 7A-F) using qualitative variables such as membrane capacitance, rheobase, resting membrane potential, neuronal excitabilities, and voltage membrane response to different injected current steps. The PCA results confirmed the lack of differentiation in the pIC among CBD-exposed progeny of both sexes (Figure 7B, E and Supplementary Figure 3B), highlighting CBD prenatal exposure as a major contributing factor to the variance in the dataset (Figure 7C, F).

### 3.2 Gestational exposure to CBD altered the excitability of pyramidal neurons in specific subregions of the IC

The observed selective changes in cellular properties (such as membrane resting potential and rheobase) strongly indicate region-specific modifications in excitability of IC principal neurons following prenatal exposure to CBD. Therefore, we investigated the intrinsic firing properties between aIC and pIC in CBD progeny of both sexes (Figure 4, Table 3, Supplementary Figure 1). Gestational CBD resulted in altered intrinsic excitability of IC pyramidal neurons in a subregion-specific manner. Within the aIC, both male and female mice exposed to CBD showed a comparable response in membrane profiles when compared to the Sham counterpart (Figure 4A, C). In contrast, pyramidal neurons in the pIC of CBD-exposed mice exhibited lower excitability compared to those of Sham group, regardless of sex (Figure 4B, D). Interestingly, when comparing the firing profiles within CBD-exposed progeny, a sex-specific effect of gestational CBD was revealed, showing lower excitability of pIC neurons in CBD-exposed females compared to those in CBD-exposed males. (Supplementary Figure 1H).

**Table 3.**
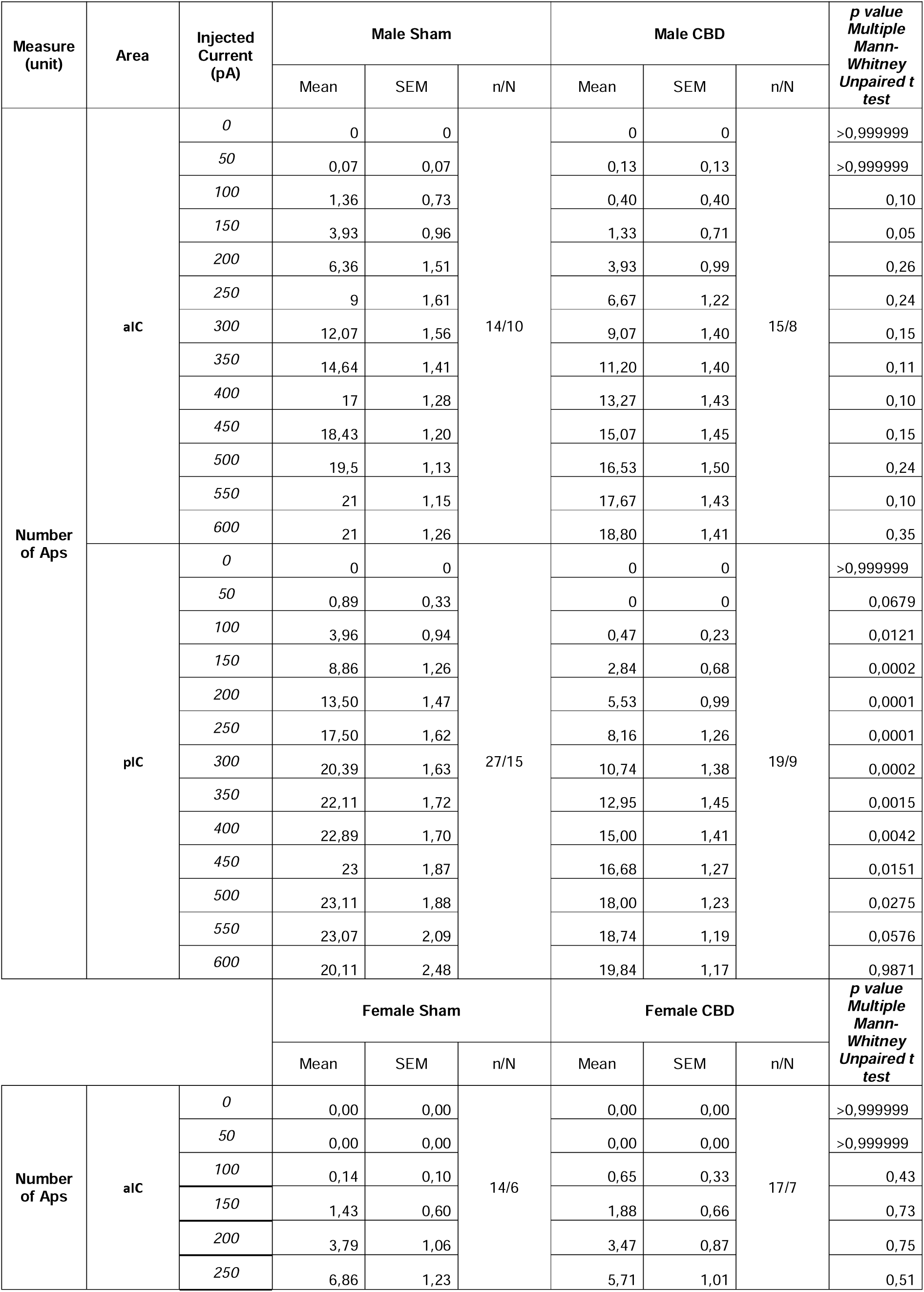
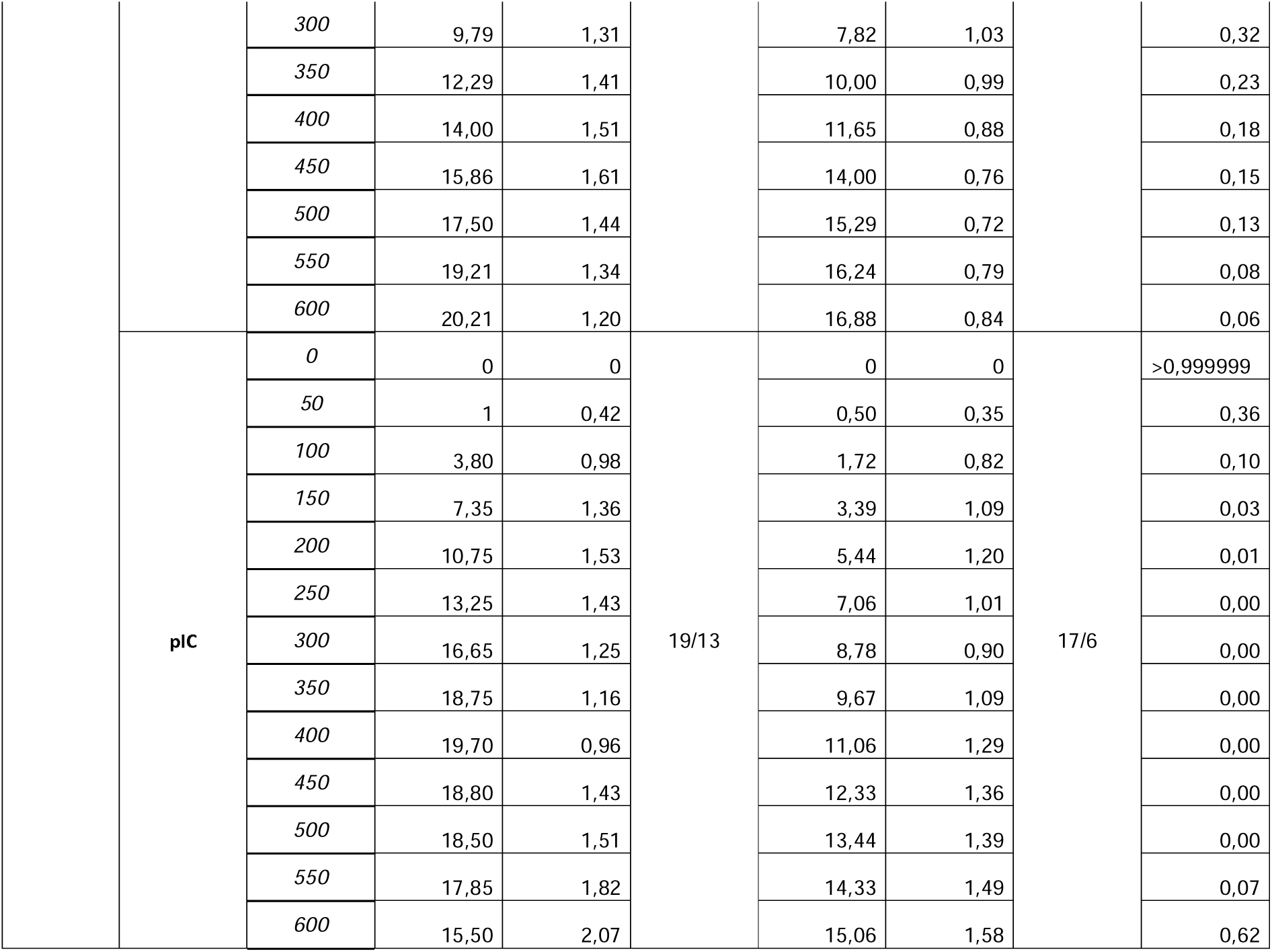
Intrinsic excitability of aIC and pIC pyramidal neurons in both sexes and IC subregions. Significance defined as p-value < 0.05.

**Figure 4.**
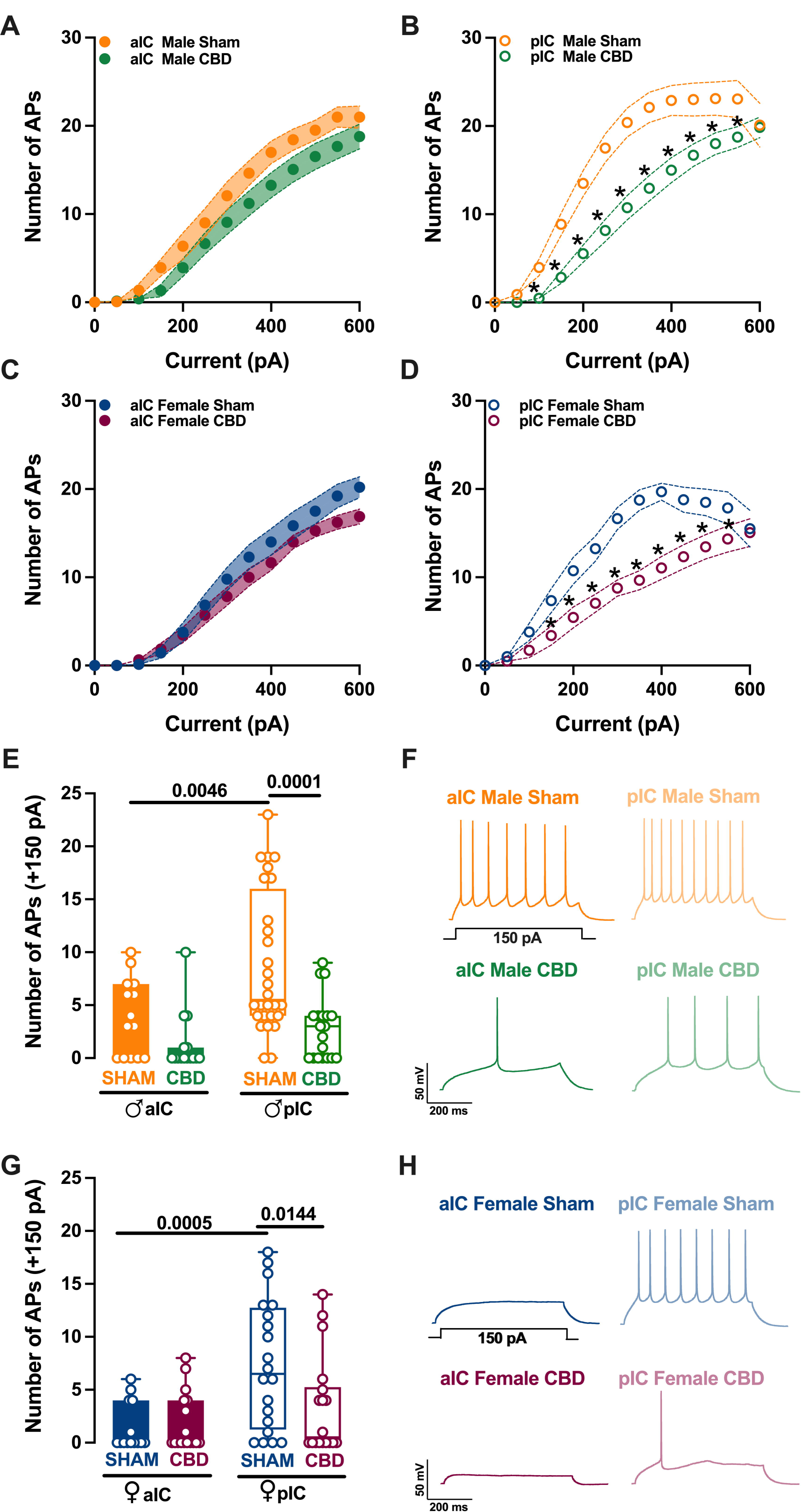
CBD prenatal exposure induces subregion-specific alterations in IC pyramidal neurons’ excitabilities, with aIC neurons being unaffected and pIC neurons showing decreased excitability. Panels **A-D** show the input-output plots for aIC and pIC pyramidal neurons, revealing that CBD prenatal exposure does not alter aIC excitability, but decreases pIC excitability, in both sexes. In contrast to Sham mice, CBD progeny exhibit a lower level of excitability in the pIC, and pyramidal neurons from pIC display less frequent firing. Panels **E-G** show the firing frequency at a physiological current step (+150 pA), demonstrating that aIC pyramidal neurons are more excitable than pIC neurons in Sham mice, but not in CBD mice. Example traces of firing patterns evoked by the injection of +150 pA depolarizing current are shown in Panels F-H for each group. Data are presented as mean ± SEM in XY plot for **(A-B-C-D)**, and as box-and-whisker plots (minimum, maximum, median) for **(E-G)**. Mann-Whitney U test was applied for **(A-B-C-D)**, and two-way ANOVA followed by Šídák’s multiple comparison test was performed for **(E-G)**. P–values < 0.05 depicted in the graph. aIC Sham male = 14/10, pIC Sham male = 27/15, aIC Sham female = 14/6, pIC Sham female = 19/13, aIC CBD male = 15/8, pIC CBD male = 19/9, aIC CBD female = 17/7, pIC CBD female = 17/6.

Within the physiological range CBD exposed animals showed a similar level of excitabilities across IC subregions (Figure 4E, G), whereas in Sham group aIC neurons fired less compared to those in pIC (Figure 4E, G), in both sexes. Accordingly with differences found in the firing profiles, within the same IC territory, pIC was less excitable in CBD compared to Sham, in both male and females (Figure 4E, G). Finally, no differences within CBD-exposed progeny were observed (Supplementary Figure 1D).

In continuity with the intrinsic properties, radar plot analysis and PCA similarly indicated a consistent absence of pIC differentiation among CBD progeny of both sexes, after prenatal CBD exposure (Figure 5 and Supplementary Figure 3).

**Figure 5.**
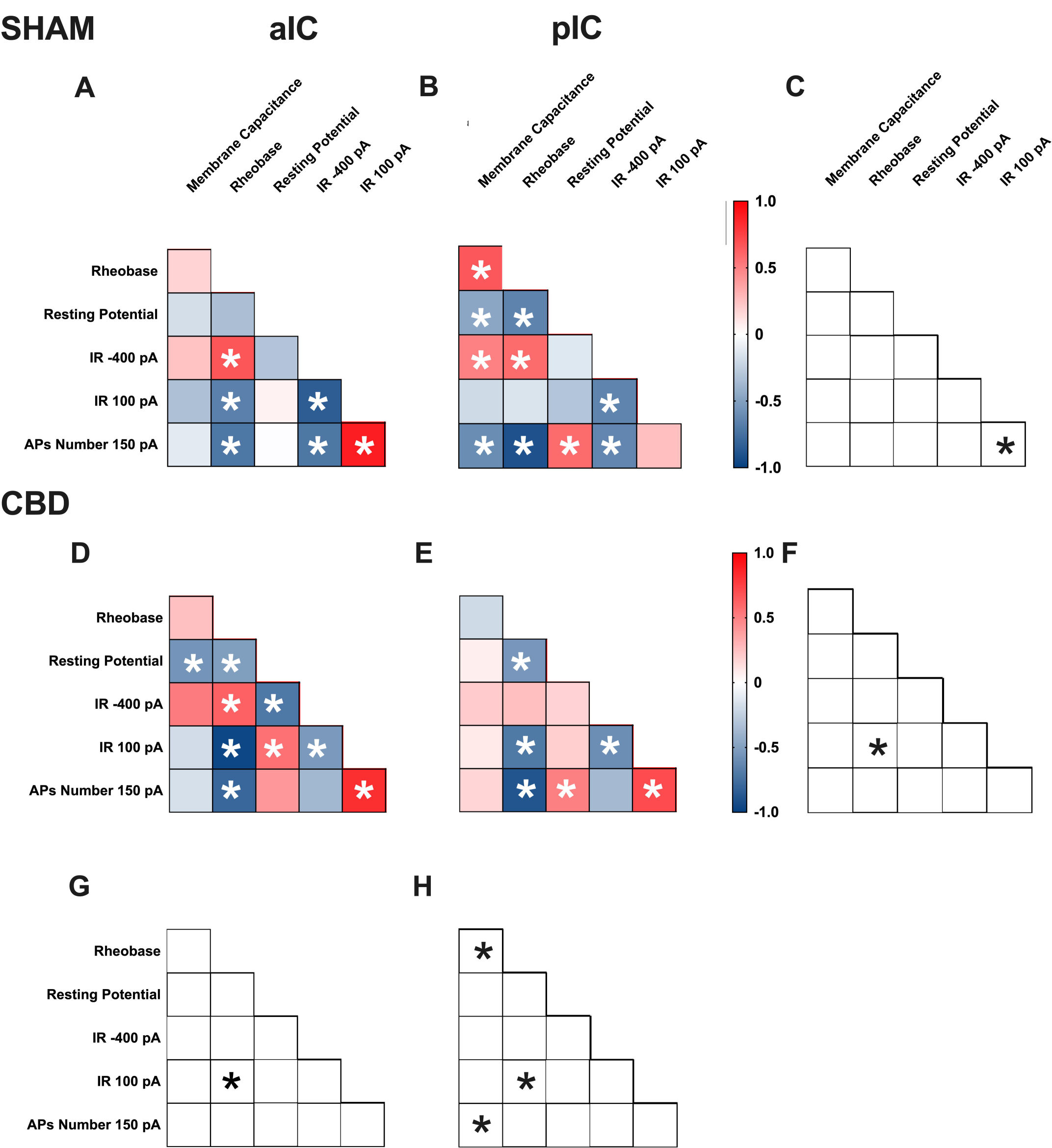
Multivariate analysis of the compound effects of prenatal CBD exposure in IC layer V pyramidal neurons of adult male offspring. **(A, B)** Heat maps of correlation profiles of electrophysiological parameters (membrane capacitance, rheobase, resting membrane potentials, APs at 150 pA, and the voltage membrane’s response to -400 and 100 pA current steps) for aIC and pIC neurons in Sham or **(D, E)** CBD exposed male mice. **(A, B, D, E)** Non-parametric Spearman correlation matrix (*r* values). Statistically significant correlations (*p*-values < 0.05) are displayed in graphs with white *. **(C, F–H)** Correlations that significantly differ (*p*-value < 0.05, Fisher’s *Z* test) are indicated with a black * by **(C, F)** subregion and **(G, H)** treatment. **(A–H)** aIC Sham male n = 14, pIC Sham male n = 27, aIC CBD male n = 15, pIC CBD male n = 19.

### 3.3 Territory-specific changes in electrophysiological correlations due to gestational CBD exposure: insights from multivariate analysis

Multivariate analysis was used to uncover complex changes in how electrophysiological parameters interact and identify patterns and correlations that may be affected like gestational CBD exposure. Specifically, we assessed how gestational exposure to CBD alters the relationships between pairs of electrophysiological features in the aIC and pIC of both sexes (Figures 5, 6, Table 4).

**Table 4.** Multivariate analysis of the compound effects of gestational CBD exposure.

| CONDITION | Measure | Membrane Capacitance | Rheobase | Resting Potential | Input Resistance -400 | Input Resistance +100 | APs number 150 pa |
| --- | --- | --- | --- | --- | --- | --- | --- |
| aIC SHAM MALE | Membrane Capacitance | 1,00 | 0,17 | -0,18 | 0,24 | -0,33 | -0,12 |
|  | Rheobase | 0,17 | 1,00 | -0,34 | 0,67 | -0,67 | -0,71 |
|  | Resting Potential | -0,18 | -0,34 | 1,00 | -0,31 | 0,05 | -0,02 |
|  | Input Resistance -400 | 0,24 | 0,67 | -0,31 | 1,00 | -0,85 | -0,72 |
|  | Input Resistance +100 | -0,33 | -0,67 | 0,05 | -0,85 | 1,00 | 0,89 |
|  | APs number 150 pa | -0,12 | -0,71 | -0,02 | -0,72 | 0,89 | 1,00 |
|  |  | Membrane Capacitance | Rheobase | Resting Potential | Input Resistance -400 | Input Resistance +100 | APs number 150 pa |
| pIC SHAM MALE | Membrane Capacitance | 1,00 | 0,66 | -0,47 | 0,50 | -0,20 | -0,60 |
|  | Rheobase | 0,66 | 1,00 | -0,65 | 0,58 | -0,15 | -0,90 |
|  | Resting Potential | -0,47 | -0,65 | 1,00 | -0,13 | -0,31 | 0,58 |
|  | Input Resistance -400 | 0,50 | 0,58 | -0,13 | 1,00 | -0,65 | -0,63 |
|  | Input Resistance +100 | -0,20 | -0,15 | -0,31 | -0,65 | 1,00 | 0,26 |
|  | APs number 150 pa | -0,60 | -0,90 | 0,58 | -0,63 | 0,26 | 1,00 |
|  |  | Membrane Capacitance | Rheobase | Resting Potential | Input Resistance -400 | Input Resistance +100 | APs number 150 pa |
| aIC SHAM FEMALE | Membrane Capacitance | 1 | -0,36 | 0,17 | 0,05 | 0,16 | 0,33 |
|  | Rheobase | -0,36 | 1,00 | 0,26 | 0,49 | -0,67 | -0,60 |
|  | Resting Potential | 0,17 | 0,26 | 1,00 | -0,06 | 0,17 | 0,12 |
|  | Input Resistance -400 | 0,05 | 0,49 | -0,06 | 1,00 | -0,78 | -0,29 |
|  | Input Resistance +100 | 0,16 | -0,67 | 0,17 | -0,78 | 1,00 | 0,55 |
|  | APs number 150 pa | 0,33 | -0,60 | 0,12 | -0,29 | 0,55 | 1,00 |
|  |  | Membrane Capacitance | Rheobase | Resting Potential | Input Resistance -400 | Input Resistance +100 | APs number 150 pa |
| pIC SHAM FEMALE | Membrane Capacitance | 1,00 | 0,44 | 0,07 | 0,39 | -0,43 | -0,51 |
|  | Rheobase | 0,44 | 1,00 | -0,36 | 0,77 | -0,51 | -0,96 |
|  | Resting Potential | 0,07 | -0,36 | 1,00 | -0,13 | -0,20 | 0,38 |
|  | Input Resistance -400 | 0,39 | 0,77 | -0,13 | 1,00 | -0,71 | -0,77 |
|  | Input Resistance +100 | -0,43 | -0,51 | -0,20 | -0,71 | 1,00 | 0,50 |
|  | APs number 150 pa | -0,51 | -0,96 | 0,38 | -0,77 | 0,50 | 1,00 |
|  |  | Membrane Capacitance | Rheobase | Resting Potential | Input Resistance -400 | Input Resistance +100 | APs number 150 pa |
| aIC CBD MALE | Membrane Capacitance | 1,00 | 0,25 | -0,56 | 0,52 | -0,19 | -0,17 |
|  | Rheobase | 0,25 | 1,00 | -0,51 | 0,62 | -0,96 | -0,80 |
|  | Resting Potential | -0,56 | -0,51 | 1,00 | -0,71 | 0,56 | 0,40 |
|  | Input Resistance -400 | 0,52 | 0,62 | -0,71 | 1,00 | -0,54 | -0,38 |
|  | **Input Resistance +100** | -0,19 | -0,96 | 0,56 | -0,54 | 1,00 | 0,82 |
|  | **APs numeber 150 pa** | -0,17 | -0,80 | 0,40 | -0,38 | 0,82 | 1,00 |
|  |  | **Membrane Capacitance** | **Rheobase** | **Resting Potential** | **Input Resistance -400** | **Input Resistance +100** | **APs numeber 150 pa** |
| **pIC CBD MALE** | **Membrane Capacitance** | 1,00 | -0,20 | 0,07 | 0,21 | 0,09 | 0,17 |
|  | **Rheobase** | -0,20 | 1,00 | -0,55 | 0,25 | -0,70 | -0,89 |
|  | **Resting Potential** | 0,07 | -0,55 | 1,00 | 0,17 | 0,19 | 0,50 |
|  | **Input Resistance -400** | 0,21 | 0,25 | 0,17 | 1,00 | -0,60 | -0,36 |
|  | **Input Resistance +100** | 0,09 | -0,70 | 0,19 | -0,60 | 1,00 | 0,71 |
|  | **APs numeber 150 pa** | 0,17 | -0,89 | 0,50 | -0,36 | 0,71 | 1,00 |
|  |  | **Membrane Capacitance** | **Rheobase** | **Resting Potential** | **Input Resistance -400** | **Input Resistance +100** | **APs numeber 150 pa** |
| **aIC CBD FEMALE** | **Membrane Capacitance** | 1,00 | -0,04 | 0,27 | 0,11 | -0,12 | -0,02 |
|  | **Rheobase** | -0,04 | 1,00 | -0,90 | 0,54 | -0,82 | -0,89 |
|  | **Resting Potential** | 0,27 | -0,90 | 1,00 | -0,55 | 0,64 | 0,72 |
|  | **Input Resistance -400** | 0,11 | 0,54 | -0,55 | 1,00 | -0,65 | -0,48 |
|  | **Input Resistance +100** | -0,12 | -0,82 | 0,64 | -0,65 | 1,00 | 0,79 |
|  | **APs numeber 150 pa** | -0,02 | -0,89 | 0,72 | -0,48 | 0,79 | 1,00 |
|  |  | **Membrane Capacitance** | **Rheobase** | **Resting Potential** | **Input Resistance -400** | **Input Resistance +100** | **APs numeber 150 pa** |
| **pIC CBD FEMALE** | **Membrane Capacitance** | 1,00 | 0,18 | -0,12 | 0,19 | -0,19 | -0,17 |
|  | **Rheobase** | 0,18 | 1,00 | -0,70 | 0,82 | -0,82 | -0,79 |
|  | **Resting Potential** | -0,12 | -0,70 | 1,00 | -0,58 | 0,62 | 0,75 |
|  | **Input Resistance -400** | 0,19 | 0,82 | -0,58 | 1,00 | -0,87 | -0,72 |
|  | **Input Resistance +100** | -0,19 | -0,82 | 0,62 | -0,87 | 1,00 | 0,87 |
|  | **APs numeber 150 pa** | -0,17 | -0,79 | 0,75 | -0,72 | 0,87 | 1,00 |

**Figure 6.**
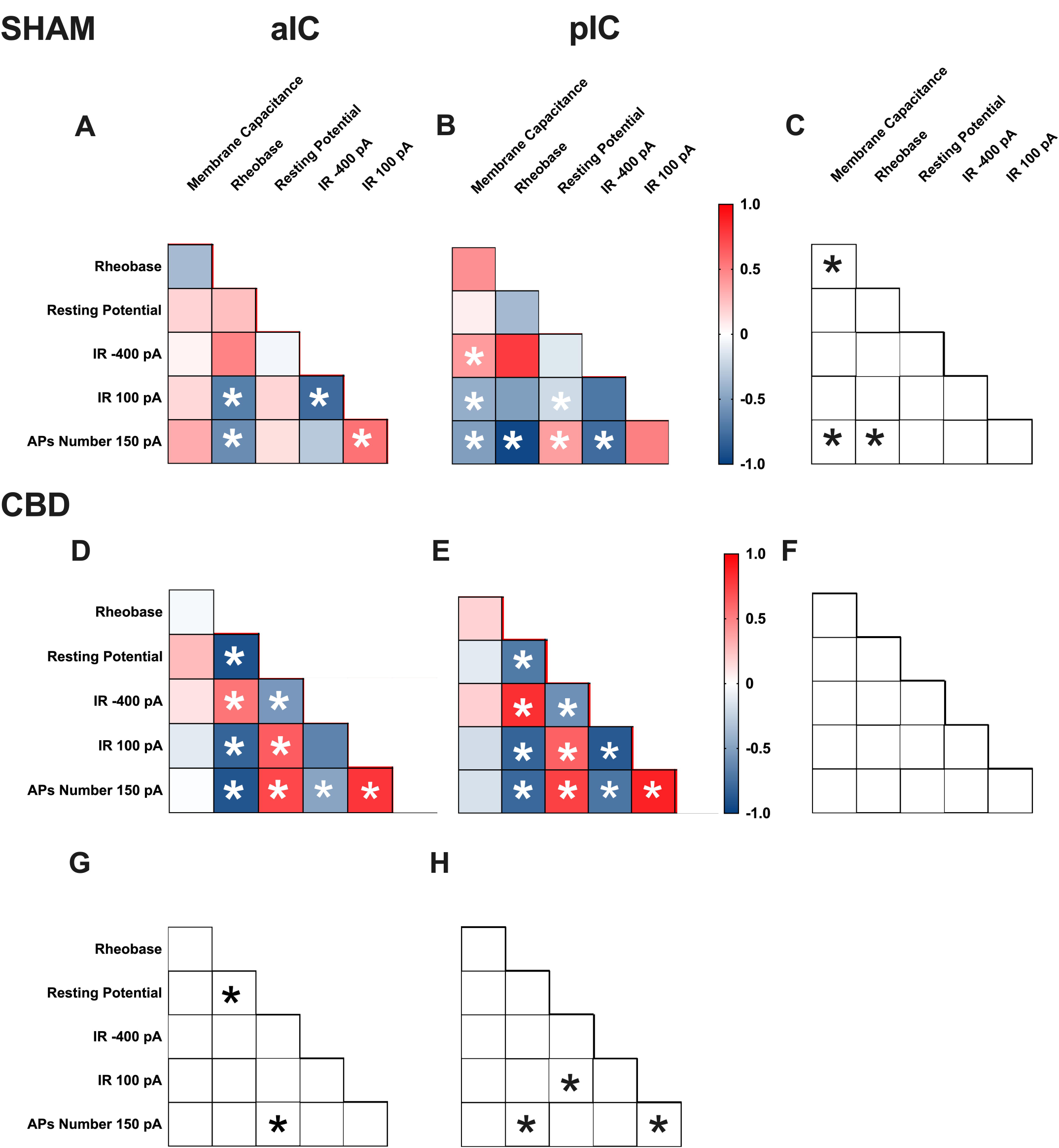
Multivariate analysis of the compound effects of prenatal CBD exposure in IC layer V pyramidal neurons of adult female offspring. **(A, B)** Heat maps of correlation profiles of electrophysiological parameters (membrane capacitance, rheobase, resting membrane potentials, APs at 150 pA, and the voltage membrane’s response to -400 and 100 pA current steps) for aIC and pIC neurons in Sham or **(D, E)** CBD exposed female mice. **(A, B, D, E)** Non-parametric Spearman correlation matrix (*r* values). Statistically significant correlations (*p*-values < 0.05) are displayed in graphs with white *. **(C, F–H)** Correlations that significantly differ (*p*-value < 0.05, Fisher’s *Z* test) are indicated with a black * by **(C, F)** subregion and **(G, H)** treatment. **(A–H)** aIC Sham female n = 14, pIC Sham female n = 21, aIC CBD female n = 17, pIC CBD female n = 18.

**Figure 7.**
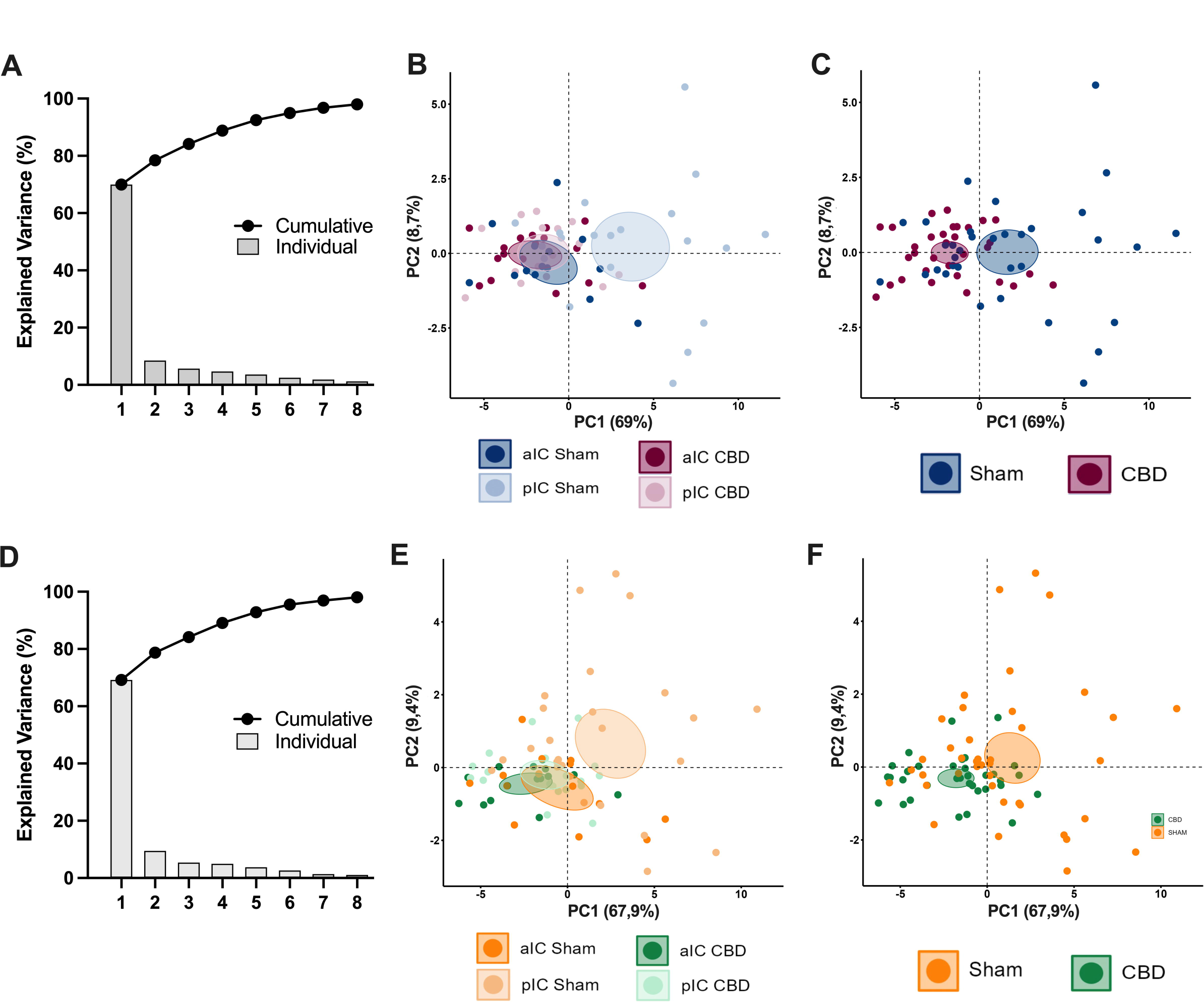
Principal component analysis (PCA) of electrophysiological properties reveals that CBD *in utero exposure* causes the lack of IC subregions differentiations in both sexes. This analysis was carried out using membrane capacitance, rheobase, resting membrane potentials, neuronal excitabilities, and the voltage membrane’s response to varying injected current steps as quantitative variables and cells as individuals. **(A, D)** Plotting the percentage of explained variance by each PC (histogram) reveals that most of the dataset’s variance is explained by PC1 (69%), PC2 (8,7%) and PC1 (67,9%), PC2 (9,4%), in female and male respectively. The cumulative percentage of explained is represented by black dots. **(B-C-E-F)** Small dots represent individuals colored according to their belonging to one the following qualitative supplementary variables: area and treatment (left), treatment (right). Each circle represents a cell plotted against its primary and secondary principal component (PC) scores. Ellipses represent the barycenter of individuals (i.e., mean) for each category, surrounded by its 95% confidence ellipses. **(B, E)** PCA showed that in Sham mice aIC and pIC largely differed in their properties, in both males and females. In contrast, IC subregions overlap in the CBD group, in both sexes. **(C, F)** This effect was driven by the CBD prenatal exposure which impacted the intrinsic properties of IC pyramidal neurons in both males and females progeny.

In Sham males, across subregions, IC principal neurons showed notable differences in the correlation between the number of action potentials (APs) fired at 150 pA and input resistance at 100 pA (Figure 5C). Conversely, male progeny exposed to CBD exhibited differences only in the correlation between input resistance at 100 pA and rheobase (Figure 5F) across IC subregions. In Sham females, differences were observed in the correlations of rheobase-membrane capacitance, the number of APs fired at 150 pA-membrane capacitance, and rheobase (Figure 6C). Interestingly, no significant correlations were found between aIC and pIC in CBD female offspring (Figure 6F). Across treatments, aIC pyramidal neurons in male progeny showed variation in the relationship between input resistance at 100 pA and rheobase (Figure 5G). In contrast, pIC neurons significantly differed in the correlations of rheobase-membrane capacitance, input resistance at 100 pA-rheobase, and the number of APs fired at 150 pA-membrane capacitance (Figure 5H). Comparing aIC of Sham and CBD female progeny revealed a significant change in the correlations between resting potential-rheobase and the number of APs fired-resting potential (Figure 6G). In pIC, changes were observed in the correlations of input resistance at 100 pA-resting potential, the number of APs fired-rheobase, and input resistance at 100 pA (Figure 6H).

This analysis, which shows complex changes in how various electrophysiological parameters interact, highlights the intricate nature of neuronal responses to gestational CBD exposure in a territory-specific manner.

### 3.4 Prenatal Cannabidiol treatment equalized excitatory and inhibitory synaptic transmission of the adult offspring in a sex- and territory-specific manner

Given the strong connections of the IC to various cortical and subcortical brain regions, we examined whether gestational CBD exposure not only affected the local circuit but also altered the connectivity of IC subregions. To assay the synaptic connectivity, we recorded both spontaneous AMPA- and GABA-mediated postsynaptic currents (sEPSCs and sIPSCs respectively) in layer V pyramidal neurons of Sham and CBD-exposed progeny of both sexes (Figures 8, 9, 10, 11, Table 5, Supplementary Figures 4 and 5). We previously showed that aIC received smaller and more frequent excitatory inputs compared to pIC in Naive male [18]. Interestingly, after prenatal CBD exposure, the difference in mean sEPSCs amplitude observed in Naive, as well as in Sham males, completely disappeared between aIC and pIC neurons in CBD-exposed males (Figure 8B). Instead, the frequency of excitatory events in aIC was higher compared to the pIC, like observed in Sham mice (Figure 8C). No differences in the kinetics of AMPA-mediated events were observed between Sham and CBD male progeny (Figure 8C-E). Contrary to that observed in males, the sEPSCs profiles in CBD females were like those of Sham females (Figure 9B-E).

**Table 5.**
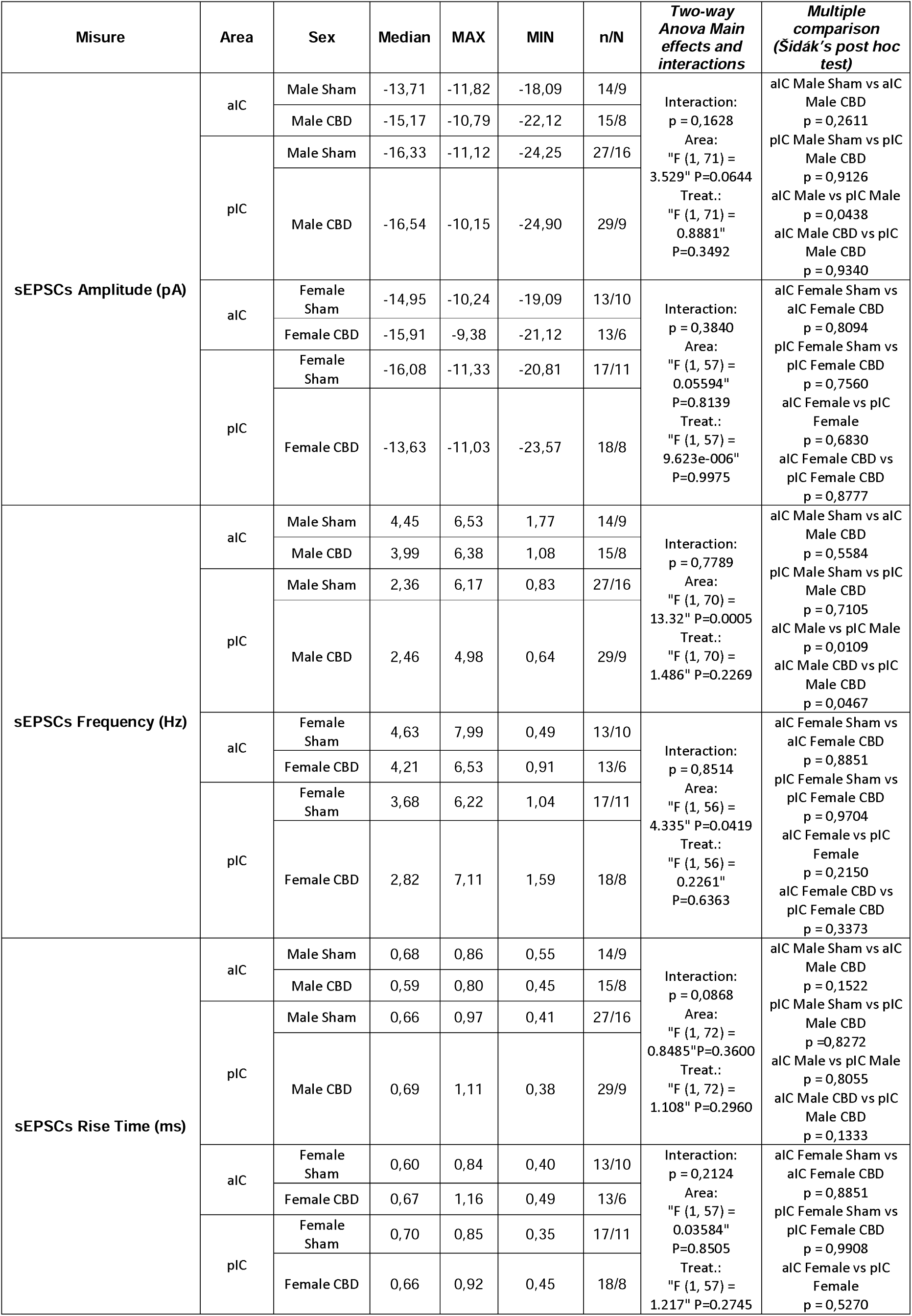
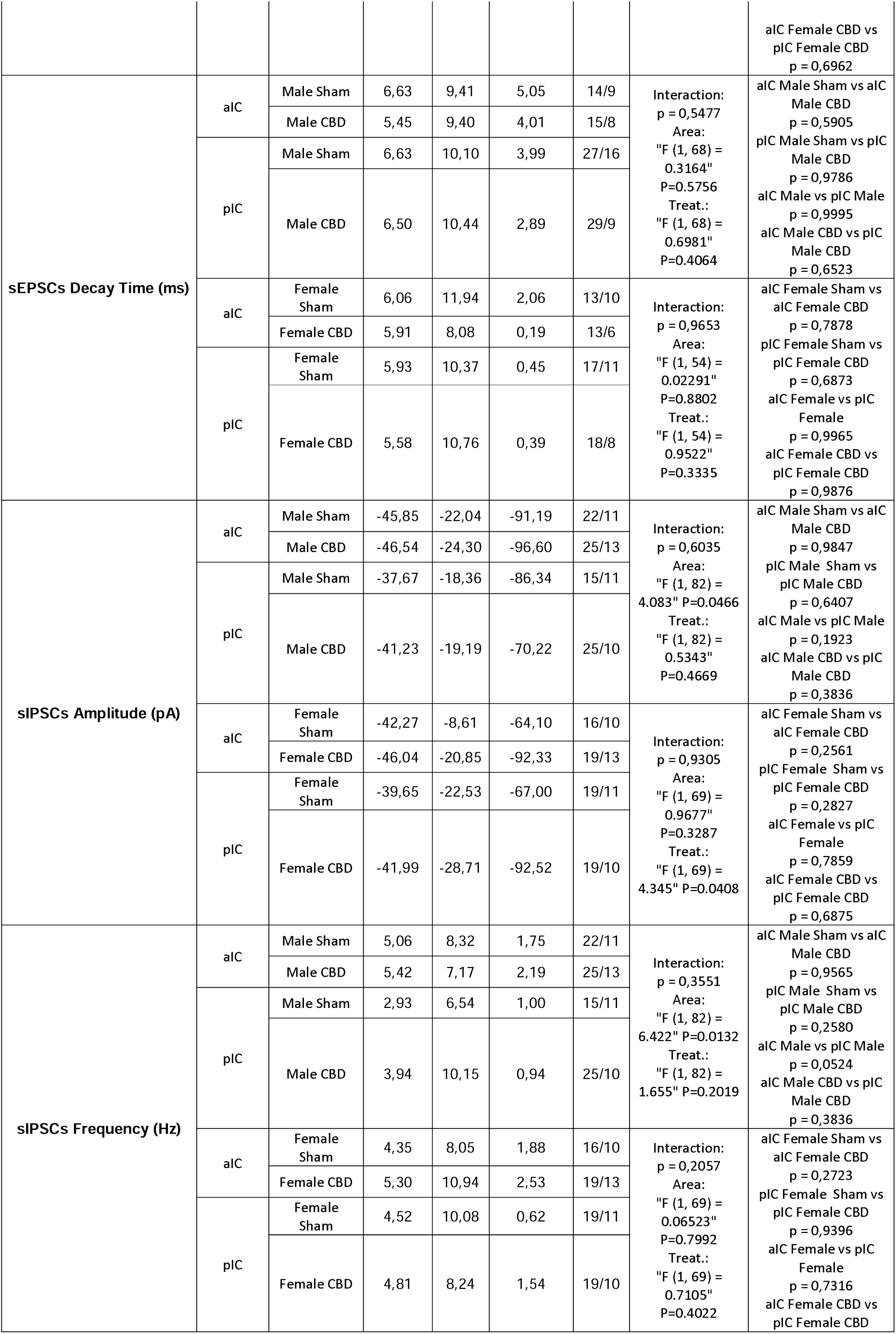
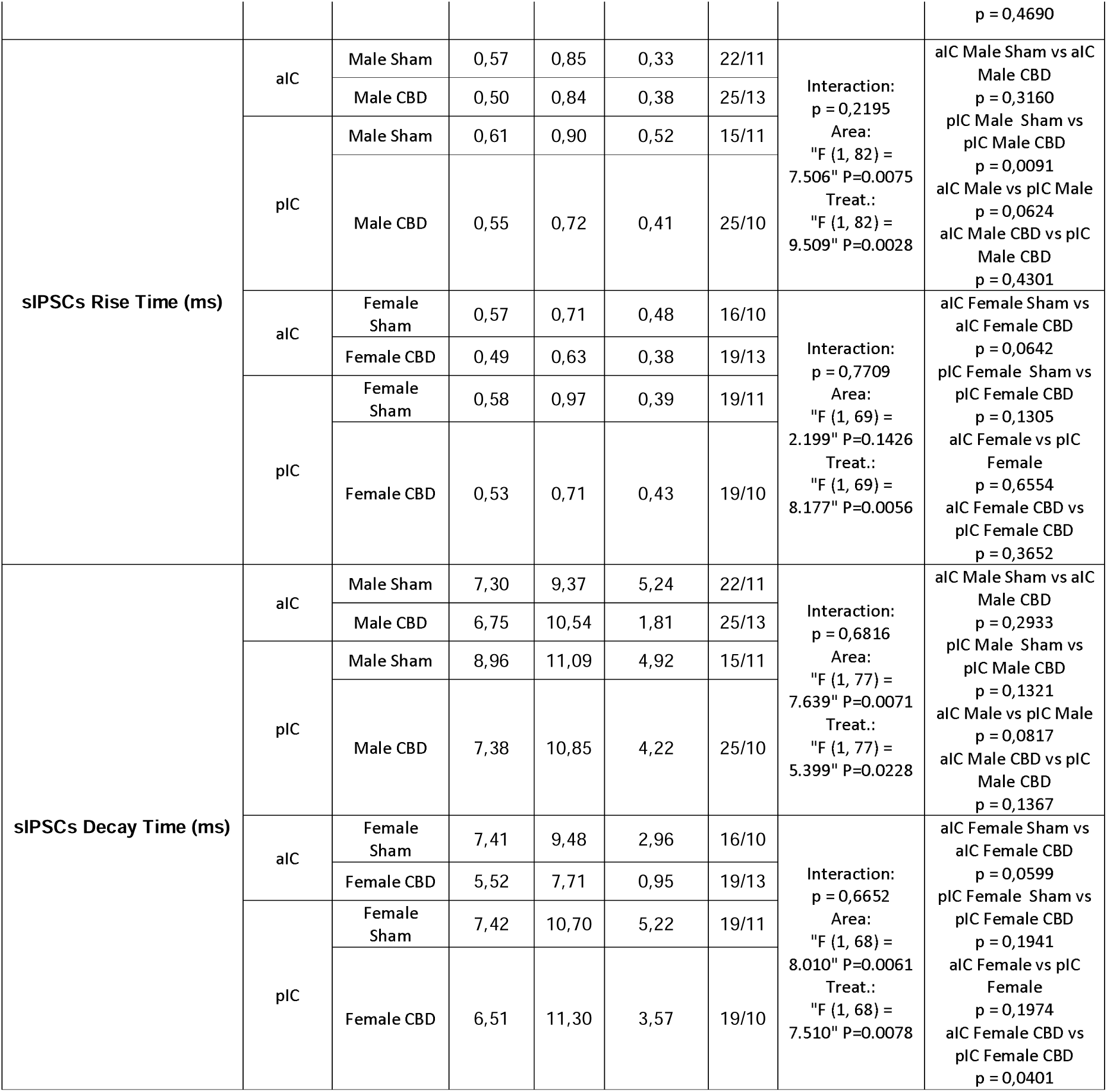
sE/IPSCs across IC subregions and treatment. Significance defined as p-value < 0.05.

**Figure 8.**
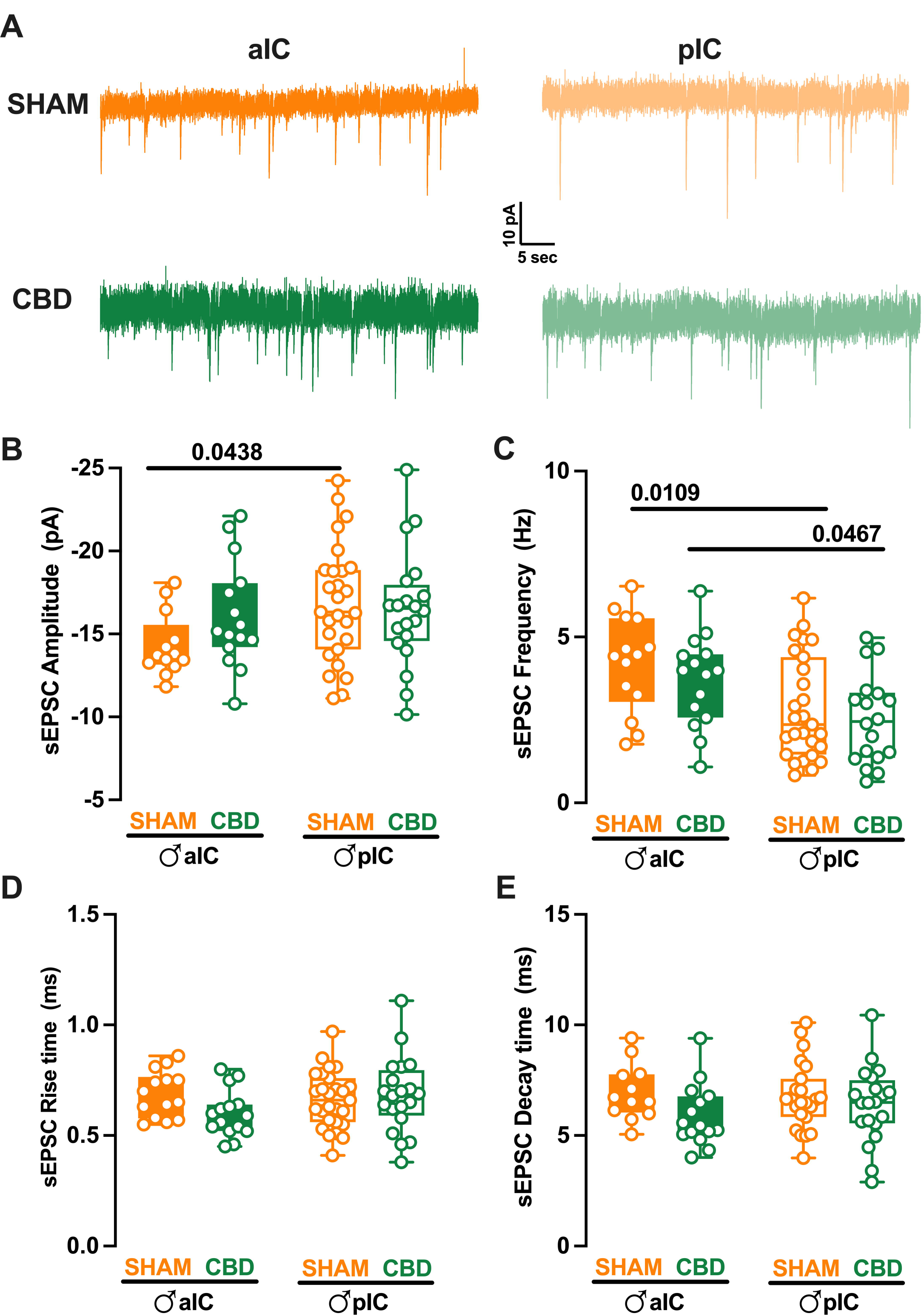
Prenatal Cannabidiol treatment affects excitatory IC synaptic transmission in a sex-specific manner, with significant effects on the amplitude of excitatory events in male progeny. Panel **A** shows representative spontaneous excitatory currents (sEPSCs) recorded at -70 mV for each group. Panels **B-E** demonstrate that prenatal CBD exposure affects the amplitude and frequency of sEPSCs in a sex-specific manner. In male progeny, CBD exposure in utero resulted in similar mean amplitudes of excitatory events in aIC and pIC, whereas in Sham mice, aIC was characterized by smaller events compared to pIC. In contrast, prenatal CBD had no effects on the frequency and kinetics of sEPSCs. Data are presented as box-and-whisker plots (minimum, maximum, median) for Panels **B-E**. Statistical analysis was performed using two-way ANOVA followed by Šídák’s multiple comparison test. *P-values < 0.05 are depicted in the graph. The sample sizes for each group were: aIC Sham male, n = 14/9; pIC Sham male, n = 27/16; aIC CBD male, n = 15/8; pIC CBD male, n = 20/9.

**Figure 9.**
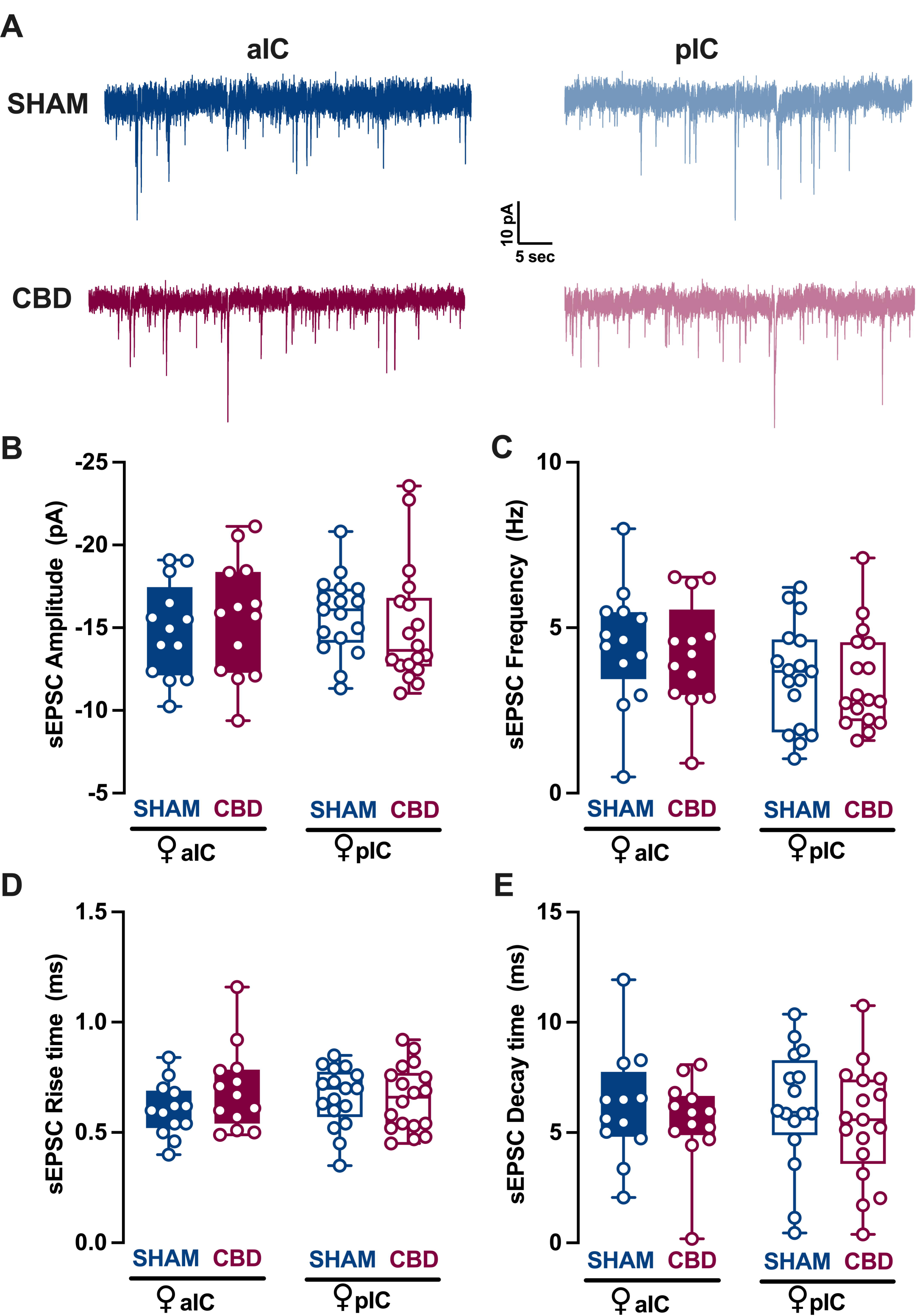
Prenatal CBD exposure does not alter excitatory synaptic transmission in principal neurons of the IC in female offspring. **(A)** Representative spontaneous excitatory postsynaptic currents (sEPSCs) recorded at -70 mV for each group. **(B-E)** Comparison of sEPSC parameters in female progeny exposed to CBD in utero versus sham-treated controls. Box-and-whisker plots (minimum, maximum, median) show that the mean amplitude **(B)**, frequency **(C)**, and kinetics **(D-E)** of sEPSCs in anterior (aIC) and posterior (pIC) IC subregions are similar between CBD-exposed and sham-treated females. Two-way ANOVA with Šídák’s multiple comparison test was used to analyze the data. No significant differences were found (p > 0.05). Sample sizes: aIC Sham female, n = 13/10; pIC Sham female, n = 17/11; aIC CBD female, n = 13/6; pIC CBD female, n = 18/8.

**Figure 10.**
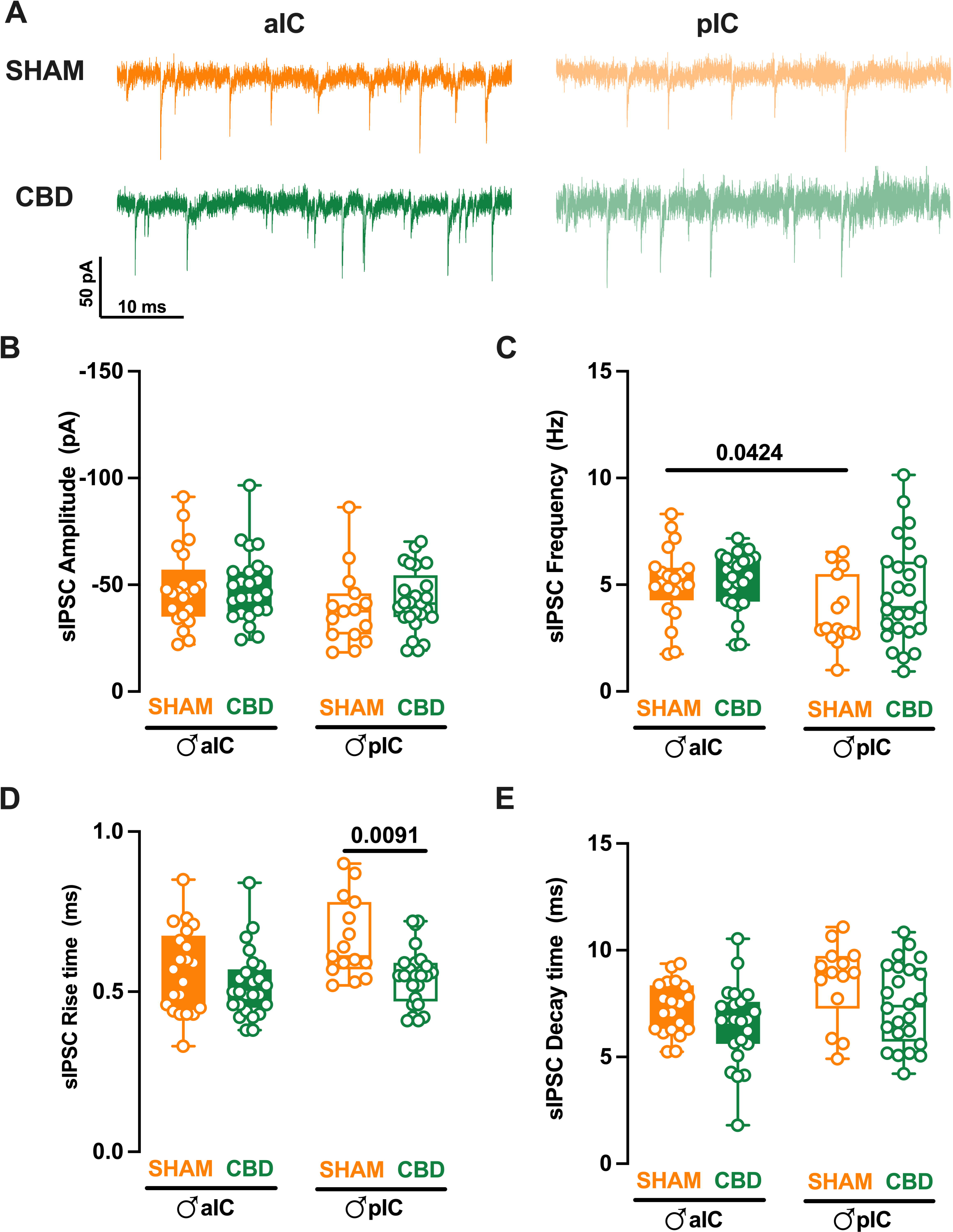
Prenatal CBD exposure normalizes IC’s inhibitory spontaneous transmission frequency and slows sIPSCs’ rise time in male offspring. **(A)** Representative spontaneous inhibitory postsynaptic currents (sIPSCs) recorded at -70 mV in anterior (aIC) and posterior (pIC) insular cortex of male Sham and CBD-exposed animals. **(B)** Quantitative analysis of sIPSC amplitude reveals no significant differences between CBD-exposed and Sham male progeny. **(C)** Gestational CBD exposure normalizes the frequency differences observed in Sham animals across IC subregions. **(D-E)** Kinetic analysis of sIPSCs shows a slower rise time in CBD-exposed males compared to Sham males in pIC, but no differences in decay time. **(B-E)** Box-and-whisker plots (minimum, maximum, median) illustrate the data. Two-way ANOVA with Šídák’s multiple comparison test was used to analyze the data. Significant differences (p < 0.05) are indicated in the graph. Sample sizes: aIC Sham male, n = 22/11; pIC Sham male, n = 15/11; aIC CBD male, n = 25/13; pIC CBD male, n = 25/10.

**Figure 11.**
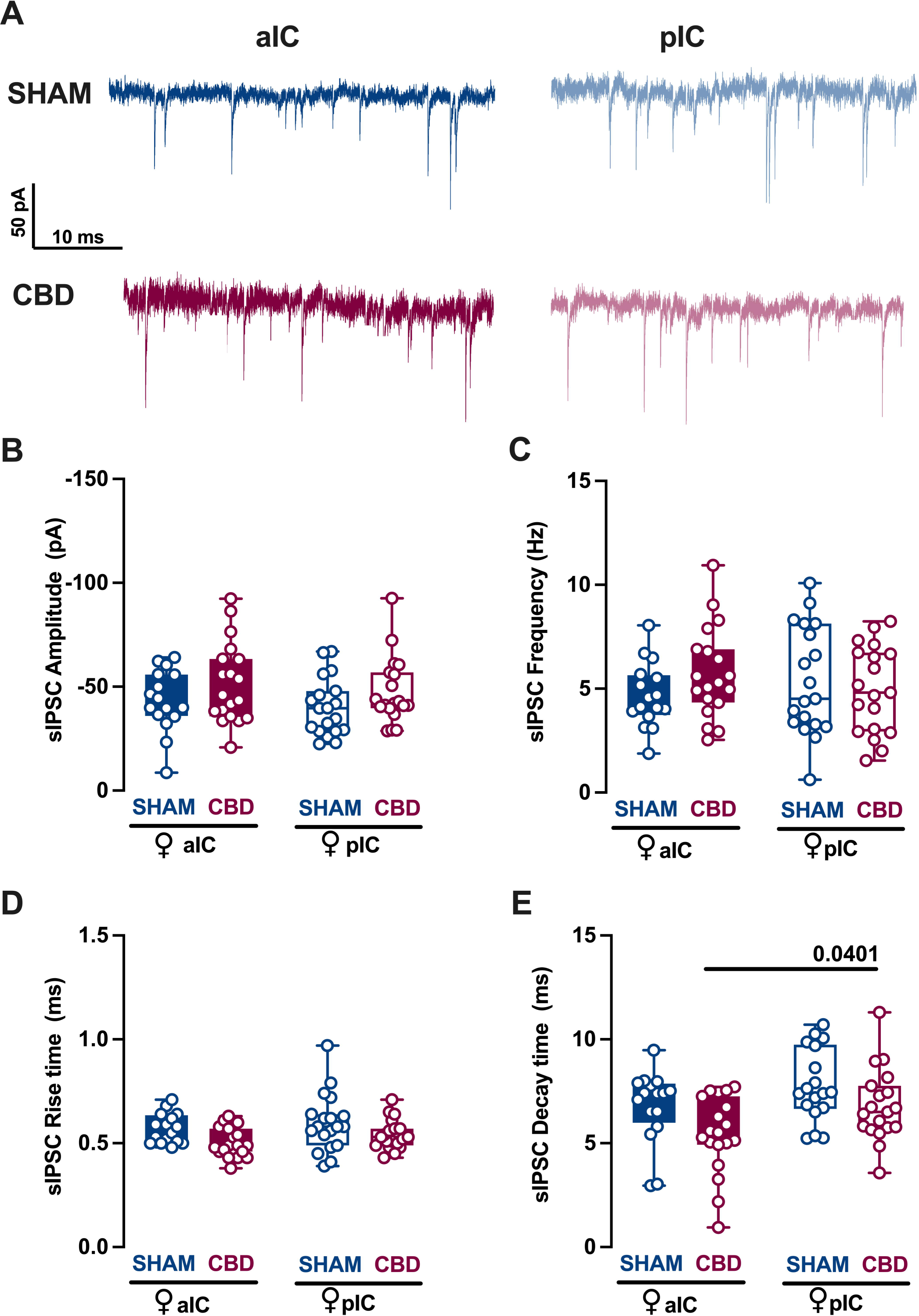
Prenatal CBD exposure has no effect on IC’s spontaneous inhibitory synaptic activity in female offspring. **(A)** Representative spontaneous inhibitory postsynaptic currents (sIPSCs) recorded at -70 mV in anterior (aIC) and posterior (pIC) insular cortex of female Sham and CBD-exposed animals. **(B-E)** Quantitative analysis reveals that inhibitory events are comparable in amplitude, frequency, and kinetics between Sham and CBD female progeny in both aIC and pIC. **(B-E)** Box-and-whisker plots (minimum, maximum, median) illustrate the data. Two-way ANOVA with Šídák’s multiple comparison test was used to analyze the data. No significant differences (p > 0.05) were found. Sample sizes: aIC Sham female, n = 16/10; pIC Sham female, n = 19/11; aIC CBD female, n = 19/13; pIC CBD female, n = 19/10.

When comparing excitatory events within the CBD progeny, a sex-specific effect of gestational CBD was revealed, showing a higher frequency of sEPSCs in the aIC compared to the pIC in males, but not in females (Supplementary Figure 4B).

We subsequently analyzed the inhibitory transmission in both Sham and CBD progeny across sexes. The mean amplitude of sIPSCs was similar between Sham and CBD males (Figure 10B). However, across IC subregions, Sham males exhibited more frequent inhibitory inputs in aIC compared to pIC (Figure 10C). Notably, prenatal exposure to CBD normalized the frequency of sIPSCs across IC subregions in CBD male (Figure 10C). Additionally, the inhibitory events in the pIC showed a slower activation in CBD males compared to Sham males (Figure 10D). On the other hand, comparing the inhibitory transmission between Sham and CBD females, no differences were observed in the mean amplitude, frequency as well as in the rise time (Figure 11B-D). However, inhibitory events in aIC were characterized by a slower decay time compared to those of pIC in only the CBD females’ group (Figure 11E). Finally, the GABA-mediated events showed a very similar profile within the CBD progeny, across sexes and IC subregions (Supplementary Figure 5A-D).

### 3.5 Prenatal Cannabidiol exposure altered the IC’s Excitatory-Inhibitory balance IC in a sex- and territory-specific manner

Prenatal exposure to CBD significantly altered excitatory and inhibitory synaptic transmission in CBD-exposed animals. Consequently, we investigated how this might impact the Excitation/Inhibition balance in the IC of CBD-exposed offspring. Thus, we quantified the total charge transferred from whole-cell recorded spontaneous AMPA-mediated EPSCs (sEPSCs; Figure 12A-B, Table 6) and GABA-mediated IPSCs (sIPSCs; Figure 12C-D, Table 6) a parameter which accounts for both frequency and amplitude of spontaneous events as previously describe [18]. Comparing our new results with those presented in Iezzi et al. [18], Figure 4, the current results demonstrate that the physiological differences in the total charge transfer of sEPSCs across IC subregions in male, completely disappear following gestational CBD exposure (Figure 12A). Conversely, no differences were observed in the total charge transfer of sEPSCs in females (Figure 12B), nor in the total charge transfer of sIPSCs across

**Table 6.**
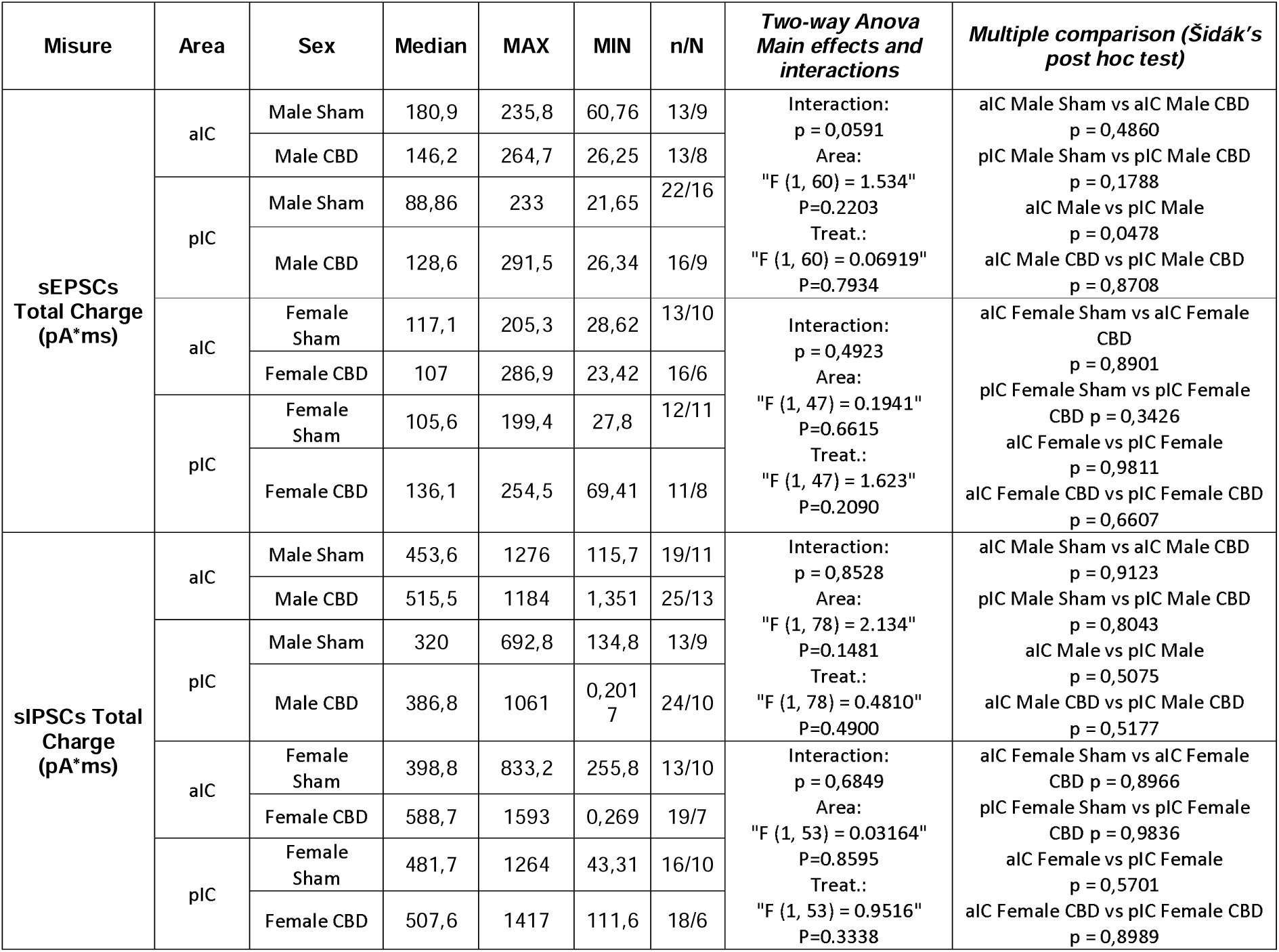
Total charge transferred by AMPA- and GABA-mediated events in aIC and pIC of both sexes and treatment. Significance defined as p-value < 0.05.

| Misure | Area | Sex | Median | MAX | MIN | n/N | Two-way Anova<br>Main effects and<br>interactions | Multiple comparison (Šidák's<br>post hoc test) |
| --- | --- | --- | --- | --- | --- | --- | --- | --- |
| sEPSCs<br>Total Charge<br>(pA*ms) | aIC | Male Sham | 180,9 | 235,8 | 60,76 | 13/9 | Interaction:<br>p = 0,0591<br>Area:<br>"F (1, 60) = 1.534"<br>P=0.2203<br>Treat.:<br>"F (1, 60) = 0.06919"<br>P=0.7934 | aIC Male Sham vs aIC Male CBD<br>p = 0,4860 |
|  |  | Male CBD | 146,2 | 264,7 | 26,25 | 13/8 |  | pIC Male Sham vs pIC Male CBD<br>p = 0,1788 |
|  | pIC | Male Sham | 88,86 | 233 | 21,65 | 22/16 |  | aIC Male vs pIC Male<br>p = 0,0478 |
|  |  | Male CBD | 128,6 | 291,5 | 26,34 | 16/9 |  | aIC Male CBD vs pIC Male CBD<br>p = 0,8708 |
|  | aIC | Female Sham | 117,1 | 205,3 | 28,62 | 13/10 | Interaction:<br>p = 0,4923<br>Area:<br>"F (1, 47) = 0.1941"<br>P=0.6615<br>Treat.:<br>"F (1, 47) = 1.623"<br>P=0.2090 | aIC Female Sham vs aIC Female<br>CBD<br>p = 0,8901 |
|  |  | Female CBD | 107 | 286,9 | 23,42 | 16/6 |  | pIC Female Sham vs pIC Female<br>CBD p = 0,3426 |
|  | pIC | Female Sham | 105,6 | 199,4 | 27,8 | 12/11 |  | aIC Female vs pIC Female<br>p = 0,9811 |
|  |  | Female CBD | 136,1 | 254,5 | 69,41 | 11/8 |  | aIC Female CBD vs pIC Female CBD<br>p = 0,6607 |
| sIPSCs Total<br>Charge<br>(pA*ms) | aIC | Male Sham | 453,6 | 1276 | 115,7 | 19/11 | Interaction:<br>p = 0,8528<br>Area:<br>"F (1, 78) = 2.134"<br>P=0.1481<br>Treat.:<br>"F (1, 78) = 0.4810"<br>P=0.4900 | aIC Male Sham vs aIC Male CBD<br>p = 0,9123 |
|  |  | Male CBD | 515,5 | 1184 | 1,351 | 25/13 |  | pIC Male Sham vs pIC Male CBD<br>p = 0,8043 |
|  | pIC | Male Sham | 320 | 692,8 | 134,8 | 13/9 |  | aIC Male vs pIC Male<br>p = 0,5075 |
|  |  | Male CBD | 386,8 | 1061 | 0,201<br>7 | 24/10 |  | aIC Male CBD vs pIC Male CBD<br>p = 0,5177 |
|  | aIC | Female Sham | 398,8 | 833,2 | 255,8 | 13/10 | Interaction:<br>p = 0,6849<br>Area:<br>"F (1, 53) = 0.03164"<br>P=0.8595<br>Treat.:<br>"F (1, 53) = 0.9516"<br>P=0.3338 | aIC Female Sham vs aIC Female<br>CBD p = 0,8966 |
|  |  | Female CBD | 588,7 | 1593 | 0,269 | 19/7 |  | pIC Female Sham vs pIC Female<br>CBD p = 0,9836 |
|  | pIC | Female Sham | 481,7 | 1264 | 43,31 | 16/10 |  | aIC Female vs pIC Female<br>p = 0,5701 |
|  |  | Female CBD | 507,6 | 1417 | 111,6 | 18/6 |  | aIC Female CBD vs pIC Female CBD<br>p = 0,8989 |

**Figure 12.**
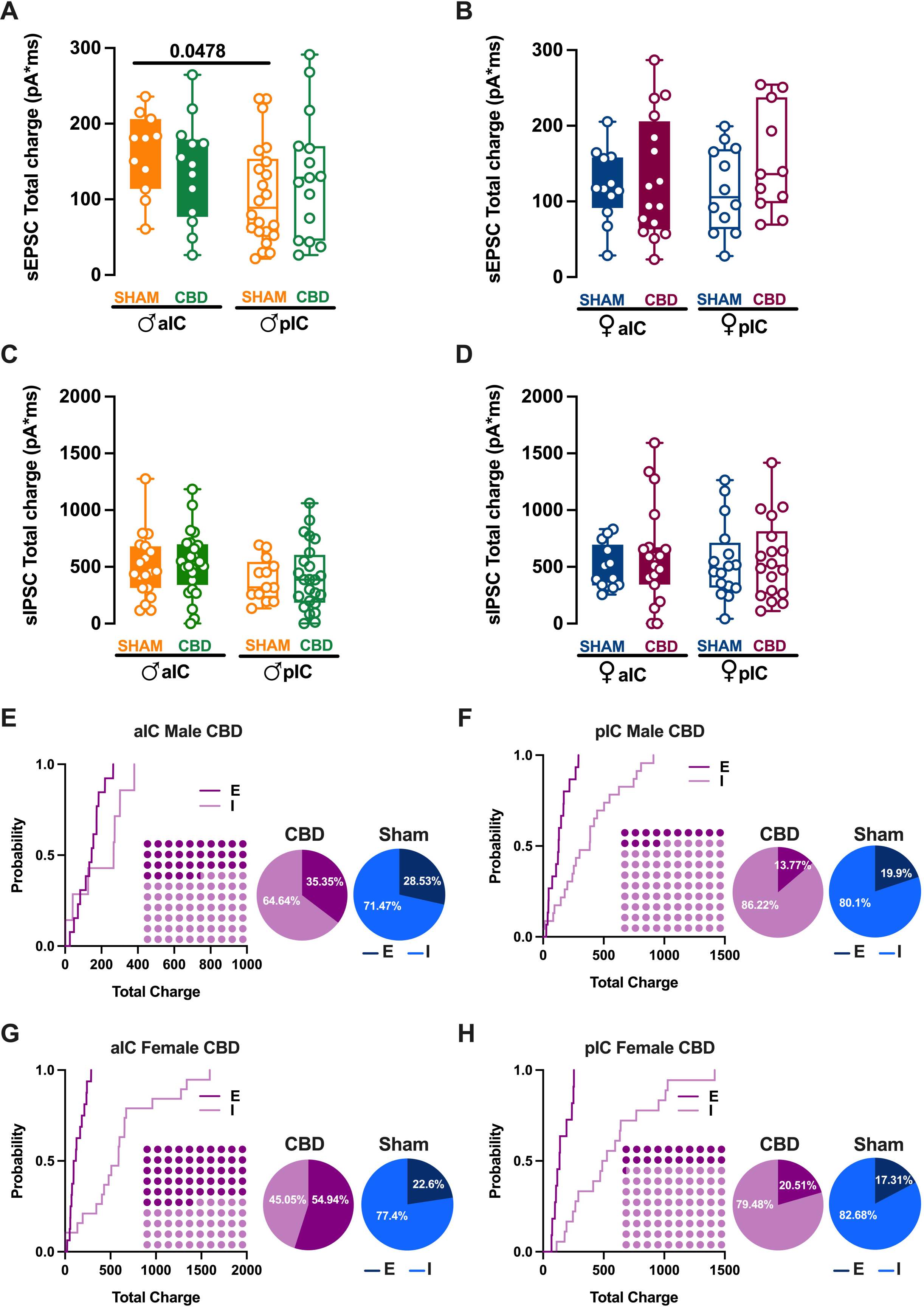
Prenatal CBD exposure altered the Excitatory/Inhibitory (E/I) balance in both IC’s subregions. **(A)** Gestational CBD exposure equalizes the total charge of AMPA-sEPSCs between aIC and pIC in male progeny, as measured over a 6-min period. **(B)** No differences in the total charge of AMPA-sEPSCs is similar across IC subregions in both Sham and CBD female progeny. **(C-D)** The total charge of GABA-sIPSCs, measured over 6-min period, shows comparable amounts of charge transferred in Sham and CBD groups of both sexes, across IC subregions. **(E-H)** Cumulative frequency distribution of sEPSCs (E) and sIPSCs (I) total charge transfer obtained from each Insular cortex within CBD male and female animals. Insets: dot plots and pie graphs (light purple/purple and light blue/blu for CBD and Sham animals respectively) showing the proportion of E versus I extrapolated at P = 0.5 from the corresponding cumulative frequency. **(A-D)** Box-and-whisker plots (minimum, maximum, median) illustrate the data, which were analyzed via two-way ANOVA followed by Šídák’s multiple comparison. Significant differences (p < 0.05) are indicated in the graphs. Sample sizes: **(A)** aIC Sham male, n = 13/9; pIC Sham male, n = 22/16; aIC CBD male, n = 13/8; pIC CBD male, n = 16/9. **(B)** aIC Sham Female, n = 13/10; pIC Sham Female, n = 12/11; aIC CBD Female, n = 16/6; pIC CBD Female, n = 11/8. **(C)** aIC Sham male, n = 19/11; pIC Sham male, n = 13/9; aIC CBD male, n = 25/13; pIC CBD male, n = 24/10. **(D)** aIC Sham Female, n = 13/10; pIC Sham Female, n = 16/10; aIC CBD Female, n = 19/7; pIC CBD Female, n = 18/6.

IC subregions and sexes (Figure 12C-D). Next, the relative distributions of sEPSCs and sIPSCs total charge transfer in CBD progeny were compared (Figure 12E-H). Despite the global prevalence of the Inhibition in the IC of both CBD progeny (Figure 12E-H, Dot plots) and Sham group (Iezzi et al., Figure 4E-H) we found differences in the distribution of Excitation and Inhibition across IC subregions and treatment in both sexes. Specifically, we observed a lower percentage of Inhibition in aIC of both CBD-treated male and females, when compared with their respective Sham counterparts (Figure 12E-G, Pie chart). In contrast, the pIC exhibited a similar percentage of Excitation and Inhibition across treatment in both sexes (Figure 12F-H, Pie chart).

## 4. Discussion

A significant proportion of pregnant women, 20.4%, report using CBD-only products, compared to 11.3% of non-pregnant women [22]. The primary reasons for CBD use among pregnant women include alleviating anxiety, depression, post-traumatic stress disorder, pain, headaches, and nausea or vomiting. Despite this trend, the effects of prenatal CBD exposure (PCE) on fetal development remain largely unknown [9, 10, 23, 24]. We investigated the impact of low-dose CBD exposure on the developing fetus. Using a mouse model, we report that PCE leads to sex-specific and region-specific changes in the functioning of the IC in adult offspring. Specifically, the results show that PCE alters the passive and active membrane properties of IC principal neurons, primarily in males, and reduces the intrinsic excitability of pyramidal neurons in both sexes. Furthermore, PCE affects synaptic transmission in the aIC of male offspring, leading to a significant shift in the excitatory/inhibitory balance within specific IC territories.

Our previous research demonstrated that in naïve mice, principal neurons in the IC exhibit distinct electrophysiological properties influenced by sex, that reflect distinct connectivity and functions within each territories of the IC [18].

Our study reveals that maternal exposure to a low dose of CBD during pregnancy abolishes the characteristic territorial differences in the IC that are normally observed in unexposed animals. In contrast to the offspring of sham-treated animals, which exhibited electrophysiological profiles similar to those of naive mice, prenatal CBD exposure was found to perturb the normal functional differentiation between aIC and pIC in a sex-specific and subregion-dependent manner, as evidenced by alterations in the active and passive membrane properties of L5 IC principal neurons. Prenatal CBD exposure was found to have a profound impact on the cellular properties of IC neurons. In male offspring, CBD exposure resulted in a uniform cellular profile across IC subregions, characterized by similar soma size, resting potential, and rheobase. Additionally, pIC neurons in male offspring were more hyperpolarized, and both male and female offspring had a higher rheobase compared to sham-treated animals. These changes led to a disruption in the normal excitability gradient between aIC and pIC. Notably, our results show that gestational CBD exposure significantly decreased the intrinsic excitability of pIC neurons in both sexes, eliminating the typical posterior-to-anterior excitability gradient. These findings collectively demonstrate that prenatal CBD exposure specifically interferes with the differentiation of IC subregions, affecting distinct cellular properties in both male and female offspring.

Multivariate analysis revealed territory-specific covariation of selected intrinsic properties following in-utero CBD exposure in both sexes. While some parameters maintained their correlations across all groups (e.g., Rheobase-AP number at 150 pA), other parameters showed different correlations across IC subregions and treatments in both sexes (e.g., APs number fired at 150 pA with Input Resistance at 100 pA and Rheobase, as well as Resting Potential with Membrane Capacitance and Rheobase). Notably, cross-correlation comparisons between IC subregions and treatments confirmed a differential impact of gestational CBD on the electrophysiological features of IC principal neurons.

The IC is unique in that it is the first cortical region to differentiate and develop in utero, a process that commences around 6 weeks after conception [25]–[28]. This early structural development lays the groundwork for the IC’s functional specialization. Moreover, the prenatal establishment of the anterior-posterior segregation of the insula, which remains relatively unchanged during infancy, may indicate that this region is particularly important for early brain functional maturation. The early appearance of an adult-like functional segregation pattern in the insula highlights its potential role in shaping early brain development Thus, PCE occurs during a critical period of IC development, as demonstrated by the alterations observed in the IC of adult offspring in this study. Although the precise mechanism of action remains unclear, it is noteworthy that CBD acts on molecular targets involved in somatosensory development during fetal life, including the serotonergic system. The serotonergic system is highly expressed during fetal development [29], [30] and in the IC [31], and alterations in 5-HT levels can lead to structural and functional reorganization of sensory afferents and intracortical microcircuitry. Consequently, disrupted development of sensory cortices may result in changes in the perception of sensory stimuli early in life, with potential long-term consequences [30].

Crucially, pIC serves as the primary recipient of dense sensory inputs from the thalamic sensory nuclei[15], which are then processed and transmitted in a posterior-to-anterior gradient through intra-insular connections to generate the final output [15]. The disruption of pIC differentiation following PCE may have significant consequences for the development of sensory and cognitive abilities. Consistent with this hypothesis, previous studies have demonstrated that PCE has a detrimental impact on the behavioral outcomes of exposed offspring, as evidenced by alterations in behavior and cognition [9], [10], [24].

In addition to altering cellular properties, PCE also disrupts the connectivity of the IC in a subregion-specific manner. Our findings reveal significant changes in excitatory and inhibitory transmission, which have a direct impact on the E/I balance within the IC. Previous research has shown that inhibitory signals dominate excitatory ones in the IC of naive mice [18], highlighting the importance of inhibitory processes in IC function. While the inhibitory tone remains consistent across IC subregions in CBD-exposed offspring, we observed a marked decrease in inhibition in the aIC of CBD-exposed progeny compared to sham-treated animals. Given the established link between GABA level modifications and various neurological and psychiatric disorders in humans and rodents [32], [33], the reduction in inhibitory tone following gestational CBD exposure may be a harbinger of negative behavioral outcomes in CBD-exposed offspring, with potential consequences for specific brain function.

In conclusion, this study provides compelling evidence of the sex- and subregion-specific impact of prenatal CBD exposure on the developmental trajectory and specialization of the IC, underscoring the potential adverse consequences of in-utero exposure to CBD. Our findings demonstrate a striking sex- and subregion-specific impact on the IC, raising concerns about the potential long-term consequences of prenatal CBD exposure on brain development and function, particularly in the context of the IC, which plays a crucial role in sensory processing, emotion regulation, and cognitive function.

## Supporting information

Supplemental Figure 1 Iezzi

Supplemental Figure 2 Iezzi

Supplemental Figure 3 Iezzi

Supplemental Figure 4 Iezzi

Supplemental Figure 5 Iezzi

## Supplementary Materials

Supplementary Figure S1: Prenatally exposure to CBD induces sex and region-specific alteration in Insular Cortex of CBD-exposed mice; Supplementary Figure S2: Prenatal CBD exposure does not change the membrane voltage response of IC pyramidal neurons; Supplementary Figure S3: Principal component analysis (PCA) of electrophysiological properties reveals the loss of IC subregions differentiation in CBD-exposed progeny; Supplementary Figure S4: Gestational CDB altered the excitatory transmission in a sex- and subregion-specific manner; Supplementary Figure S5: The comparison of in utero Cannabidiol on the inhibitory transmission of CBD-exposed males and females.

## Authors’ contributions

DI: Conceptualization, Data curation, Formal analysis, Validation, Writing—original draft, review and editing. AC: Formal analysis. JPS: Statistical analysis. PC: Conceptualization, Supervision. OJM: Conceptualization, Supervision, Funding acquisition, Methodology, Project administration, Writing—original draft, review, and editing.

## Ethics declarations

### Ethics approval and consent to participate

Animals were treated in compliance with the European Communities Council Directive (86/609/EEC) and the United States National Institutes of Health Guide for the care and use of laboratory animals. All procedures using experimental animals were approved by Aix-Marseille University Institutional Animal Care & Use Committee.

## Consent for publication

All authors read and approved the final manuscript for publication.

## Availability of data and materials

All data reported in this paper will be shared by the lead contact upon request. This paper does not report original code. Any additional information required to reanalyze the data reported in this paper is available from the lead contact upon request.

## Acknowledgements

The authors are grateful to the Chavis-Manzoni team members for helpful discussions.

## Supplementary figure legends

**Supplemental Figure 1: Prenatally exposure to CBD induces sex and region-specific alteration in Insular Cortex of CBD-exposed mice. (A-B)** Quantitative analysis of passive and active membrane properties of CBD-exposed male and females unveiled that across IC subregions, pyramidal neurons of aIC are bigger and more hyperpolarized compared to those in pIC only female progeny, but not in male. **(C-D)** Pyramidal neurons of both IC did not differ in the rheobase as well as in the numbers of APs evoked at +150 pA. **(E-F)** The number of evoked action potentials in response to depolarizing current steps were similar across IC subregions in both sexes. **(G-H)** In contrast, within the same area, the pIC of CBD female showed a lower excitability compared to the pIC of CDB male. Data are presented as box-and-whisker plots (minimum, maximum, median) for **(A-B-C-D)** and as mean ± SEM in XY plot for **(E-F-G-H)**. Two-way ANOVA followed by Šídák’s multiple comparison test was performed for **(A-B-C-D)**, while Mann-Whitney U test was applied for **(E-F-G-H)**. P–values < 0.05 depicted in the graph. aIC CBD male = 15/8, pIC CBD male = 19/9, aIC CBD female = 17/7, pIC CBD female = 17/6.

**Supplemental Figure 2: Prenatal CBD exposure does not change the membrane voltage response of IC pyramidal neurons. (A-D)** In response to current injections of 50 pA from -400 pA to +50 pA the membrane profile of pyramidal neurons was similar across and within IC sub regions in both sexes. Data are presented as mean ± CI in XY plots for **(A-D)**. A Mann-Whitney U test was used for the statistical analysis, and *p-value < 0.05 was considered significant. aIC CBD male = 15/8, pIC CBD male = 20/9, aIC CBD Female = 13/6, pIC CBD Female = 18/8.

**Supplemental Figure 3: Principal component analysis shows that gestational CBD abolished IC subregions differentiation in both sexes.** Data were analyzed via PCA with membrane capacitance, rheobase, resting membrane potentials, neuronal excitabilities, and the voltage membrane’s response to varying injected current steps as quantitative variables and cells as individuals. **(A)** Plotting the percentage of explained variance by each PC (histogram) reveals that most of the dataset’s variance is explained by PC1 (64,9%) and PC2 (8%). The cumulative percentage of explained is represented by black dots. **(B-C)** Small dots represent individuals colored according to their belonging to one the following qualitative supplementary variables: sex and treatment(left) and sex (right). Each circle represents a cell plotted against its primary and secondary principal component (PC) scores. Ellipses represent the barycenter of individuals (i.e., mean) for each category, surrounded by its 95% confidence ellipses. **(B)** Group analysis revealed that this effect is driven by a loss of IC subregion differentiation following CBD in utero exposure. **(C)** PCA showed that in CBD males and females largely overlap. aIC CBD male = 15/8, pIC CBD male = 19/9, aIC CBD female = 17/7, pIC CBD female = 17/6.

**Supplementary Figure 4. Gestational CDB altered the excitatory transmission in a sex- and subregion-specific manner.** A quantitative analysis of mean amplitude and frequency in relation to area and sex demonstrated a similar amplitude **(A)** but lower frequency **(B)** of pIC excitatory events compared to aIC events in adult males only. **(C-D)** The kinetics of sEPSC were consistent across both areas and sexes. **(A-D**) Individual neurons are represented by single dots. Data are displayed as box and whisker plots (min., max., median). A two-way ANOVA followed by a Šídák multiple comparison test was used for data analysis. P-values <0.05 are indicated in the graphs. aIC male is represented as 9/14 in dark orange, pIC male as 16/27 in dark blue, aIC female as 6/13 in light orange, and pIC female as 12/17 in light blue.

**Supplementary Figure 5. The comparison of in utero Cannabidiol on the inhibitory transmission of CBD-exposed males and females. (A-D)** No differences in the mean amplitude, frequency and the kinetics of sIPSCs in CBD progeny of both sexes were observed. **(A-D)** Data are displayed as box and whisker plots (min., max., median). **(A-D)** A two-way ANOVA followed by a Šídák multiple comparison test was used for data analysis. P-values <0.05 are indicated in the graphs. aIC CBD male = 25/13, pIC CBD male = 25/10, aIC CBD female = 19/13, pIC CBD female 19/10.

## References

[1] A. C. E. Rokeby, B. V Natale, and D. R. C. Natale, “Cannabinoids and the placenta: Receptors, signaling and outcomes", Placenta, 2023;135:51–61

[2] R. Abuhasira, L. Shbiro, and Y. Landschaft, “Medical use of cannabis and cannabinoids containing products - Regulations in Europe and North America,” Eur. J. Intern. Med., 2018, vol. 49, pp. 2–6.

[3] S. Sarrafpour et al., “Considerations and Implications of Cannabidiol Use During Pregnancy,” Curr. Pain Headache Rep., 2020, 10;24(7):38.

[4] V. Feinshtein, Z. Ben-Zvi, T. Eshkoli, B. Sheizaf, E. Sheiner, and G. Holcberg, “Cannabidiol enhances xenobiotic permeability through the human placental barrier by direct inhibition of breast cancer resistance protein: an ex vivo study,” Am J Obstet Gynecol. 2013 Dec;209(6):573.e1-573.e15.

[5] P. Alves, C. Amaral, N. Teixeira, and G. Correia-da-Silva, “Cannabidiol disrupts apoptosis, autophagy and invasion processes of placental trophoblasts,” Arch. Toxicol., vol. 95, no. 10, pp. 3393–3406, 2021.

[6] M. J. Moss et al., “Cannabis use and measurement of cannabinoids in plasma and breast milk of breastfeeding mothers,” Pediatr Res 90, 861–868 (2021)

[7] E. Bekman, T. Barata, C. Miranda, S. Vaz, C. Ferreira, and A. Quintas, “Might synthetic cannabinoids influence neural differentiation?,” Ann. Med., vol. 53, no. sup1, Apr. 2021.

[8] W. Ochiai et al., “Maternal and fetal pharmacokinetic analysis of cannabidiol during pregnancy in mice,” Drug Metab. Dispos., vol. 49, no. 4, pp. 337–343, 2021

[9] D. Iezzi, A. Caceres-Rodriguez, P. Chavis, and O. J. J. Manzoni, “In utero exposure to cannabidiol disrupts select early-life behaviors in a sex-specific manner,” Transl. Psychiatry, vol. 12, no. 1, 2022, doi: 10.1038/s41398-022-02271-8.

[10] I. de S. Maciel et al., “Perinatal CBD or THC Exposure Results in Lasting Resistance to Fluoxetine in the Forced Swim Test: Reversal by Fatty Acid Amide Hydrolase Inhibition,” Cannabis Cannabinoid Res., vol. X, no. X, pp. 1–10, 2021, doi: 10.1089/can.2021.0015.

[11] N. Gogolla, “The insular cortex,” Curr. Biol., vol. 27, no. 12, pp. R580–R586, 2017, doi: 10.1016/j.cub.2017.05.010.

[12] C. Lamm and T. Singer, “The role of anterior insular cortex in social emotions,” Brain Struct. Funct. 2010 2145, vol. 214, no. 5, pp. 579–591, Apr. 2010, doi: 10.1007/S00429-010-0251-3.

[13] E. T. Rolls, “Limbic Structures, Emotion, and Memory,” Curated Ref. Collect. Neurosci. Biobehav. Psychol., Jan. 2017, doi: 10.1016/B978-0-12-809324-5.06857-7.

[14] C. Ibrahim, B. Le Foll, and L. French, “Transcriptomic Characterization of the Human Insular Cortex and Claustrum,” Front. Neuroanat., vol. 13, Nov. 2019, doi: 10.3389/FNANA.2019.00094.

[15] D. A. Gehrlach et al., “A whole-brain connectivity map of mouse insular cortex,” Elife, vol. 9, pp. 1–78, 2020, doi: 10.7554/ELIFE.55585.

[16] M. Nagai, K. Kishi, and S. Kato, “Insular cortex and neuropsychiatric disorders: a review of recent literature,” Eur. Psychiatry, vol. 22, no. 6, pp. 387–394, Sep. 2007, doi: 10.1016/J.EURPSY.2007.02.006.

[17] P. Pressman and H. J. Rosen, Disorders of Frontal Lobe Function. Elsevier Inc., 2015.

[18] D. Iezzi, A. Cáceres-Rodríguez, B. Strauss, P. Chavis, and O. J. Manzoni, “Sexual differences in neuronal and synaptic properties across subregions of the mouse insular cortex,” Biol. Sex Differ., vol. 15, no. 1, pp. 1–21, Dec. 2024, doi: 10.1186/S13293-024-00593-4/TABLES/7.

[19] B. Diedenhofen and J. Musch, “cocor: A Comprehensive Solution for the Statistical Comparison of Correlations,” PLoS One, vol. 10, no. 4, p. e0121945, Apr. 2015, doi: 10.1371/JOURNAL.PONE.0121945.

[20] “R: The R Project for Statistical Computing.” https://www.r-project.org/ (accessed Jun. 17, 2024).

[21] M. Doi, N. Usui, and S. Shimada, “Prenatal Environment and Neurodevelopmental Disorders,” Front. Endocrinol. (Lausanne)., vol. 13, p. 860110, Mar. 2022, doi: 10.3389/FENDO.2022.860110/BIBTEX.

[22] D. Bhatia, S. Battula, S. Mikulich-Gilbertson, J. Sakai, and D. Hammond, “Cannabidiol-Only Product Use in Pregnancy in the United States and Canada: Findings From the International Cannabis Policy Study,” Obstet. Gynecol., May 2024, doi: 10.1097/AOG.0000000000005603.

[23] K. Lee et al., “Cannabidiol Exposure During Gestation Leads to Adverse Cardiac Outcomes Early in Postnatal Life in Male Rat Offspring,” Cannabis Cannabinoid Res., 2024, doi: 10.1089/CAN.2023.0213/SUPPL_FILE/SUPPL_TABLES5.PDF.

[24] K. S. Swenson et al., “Fetal cannabidiol (CBD) exposure alters thermal pain sensitivity, problem-solving, and prefrontal cortex excitability,” Mol. Psychiatry 2023 288, vol. 28, no. 8, pp. 3397–3413, Jul. 2023, doi: 10.1038/s41380-023-02130-y.

[25] M. Catala, “Development of the Central Nervous System,” *Textb*. Pediatr. Neurosurg., pp. 1–99, 2019, doi: 10.1007/978-3-319-31512-6_1-1.

[26] A. Afif, R. Bouvier, A. Buenerd, J. Trouillas, and P. Mertens, “Development of the human fetal insular cortex: study of the gyration from 13 to 28 gestational weeks,” Brain Struct. Funct., vol. 212, no. 3–4, pp. 335–346, Dec. 2007, doi: 10.1007/S00429-007-0161-1.

[27] M. Y. S. Kalani, M. A. Kalani, R. Gwinn, B. Keogh, and V. C. K. Tse, “Embryological development of the human insula and its implications for the spread and resection of insular gliomas,” Neurosurg. Focus, vol. 27, no. 2, 2009, doi: 10.3171/2009.5.FOCUS0997.

[28] H. C. Evrard and A. D. (Bud) Craig, “Insular Cortex,” Brain Mapp. An Encycl. Ref., vol. 2, pp. 387–393, May 2023, doi: 10.1016/B978-0-12-397025-1.00237-2.

[29] S. Brummelte, E. Mc Glanaghy, A. Bonnin, and T. F. Oberlander, “DEVELOPMENTAL CHANGES IN SEROTONIN SIGNALING: IMPLICATIONS FOR EARLY BRAIN FUNCTION, BEHAVIOR AND ADAPTATION,” Neuroscience, vol. 342, p. 212, Feb. 2017, doi: 10.1016/J.NEUROSCIENCE.2016.02.037.

[30] S. I. Hanswijk et al., “Gestational Factors throughout Fetal Neurodevelopment: The Serotonin Link,” Int. J. Mol. Sci., vol. 21, no. 16, pp. 1–42, Aug. 2020, doi: 10.3390/IJMS21165850.

[31] A. Ju, B. Fernandez-Arroyo, Y. Wu, D. Jacky, and A. Beyeler, “Expression of serotonin 1A and 2A receptors in molecular- and projection-defined neurons of the mouse insular cortex,” Mol. Brain, vol. 13, no. 1, Jun. 2020, doi: 10.1186/S13041-020-00605-5.

[32] C. J. Watson, “Insular balance of glutamatergic and GABAergic signaling modulates pain processing,” Pain, vol. 157, no. 10, pp. 2194–2207, Jun. 2016, doi: 10.1097/J.PAIN.0000000000000615.

[33] I. M. Rosso, M. R. Weiner, D. J. Crowley, M. M. Silveri, S. L. Rauch, and J. E. Jensen, “Insula and anterior cingulate GABA levels in post-traumatic stress disorder: Preliminary findings using magnetic resonance spectroscopy,” Depress. Anxiety, vol. 31, no. 2, p. 115, Feb. 2014, doi: 10.1002/DA.22155.

