## Supplementary figures and images for "Gestational CBD shapes insular cortex in adulthood"

### Supplemental Figure 1 Iezzi

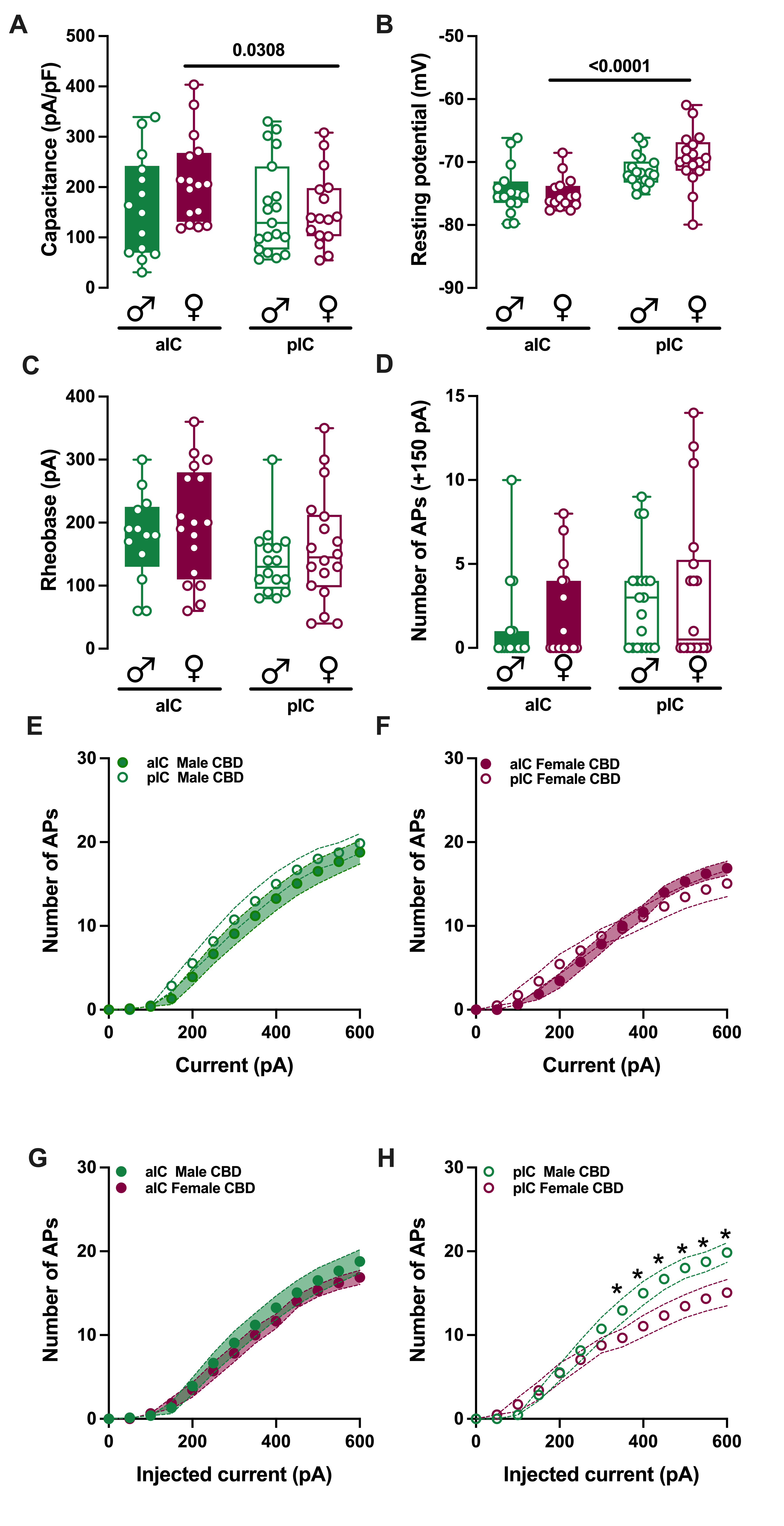

### Supplemental Figure 2 Iezzi

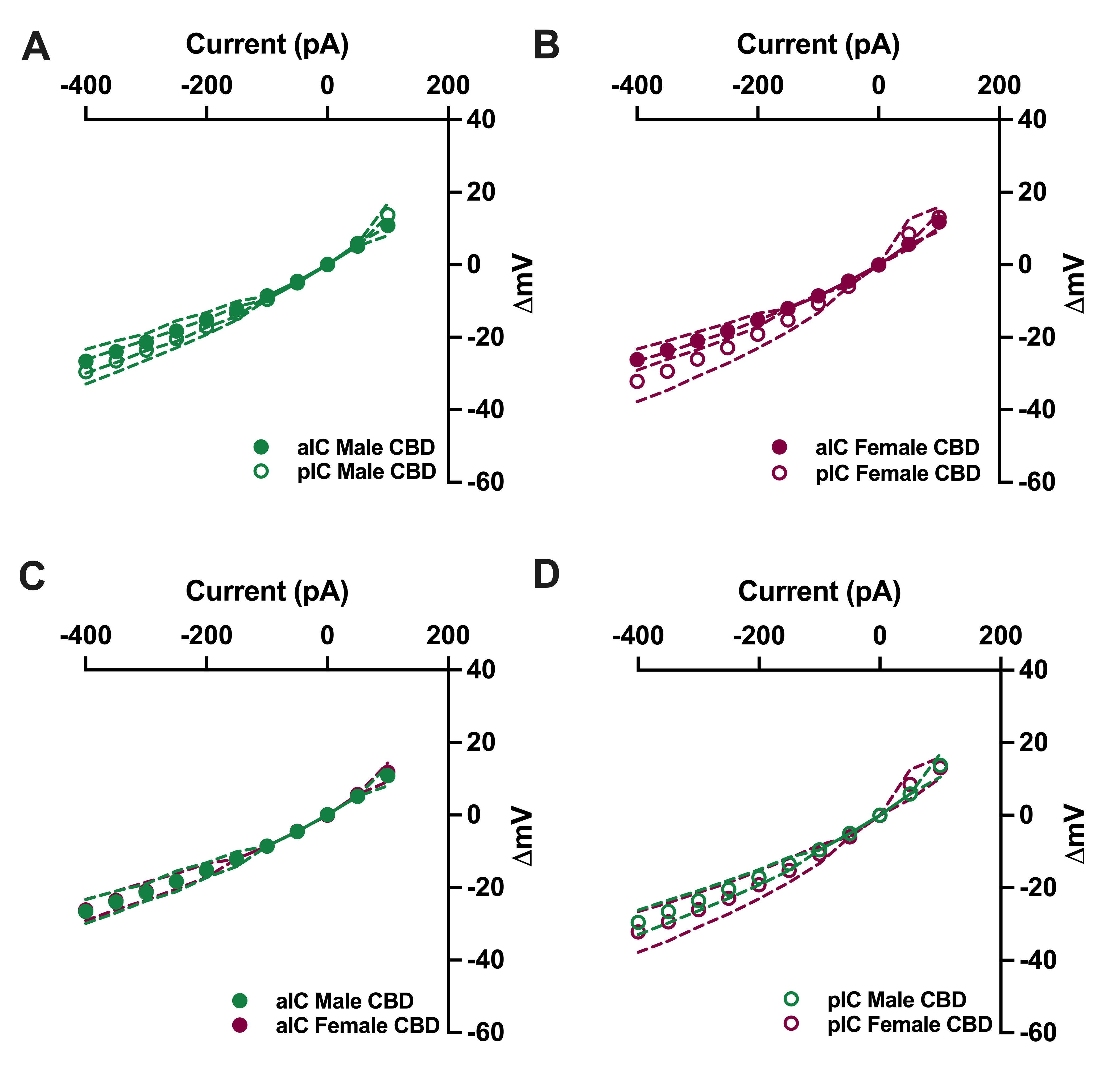

### Supplemental Figure 3 Iezzi

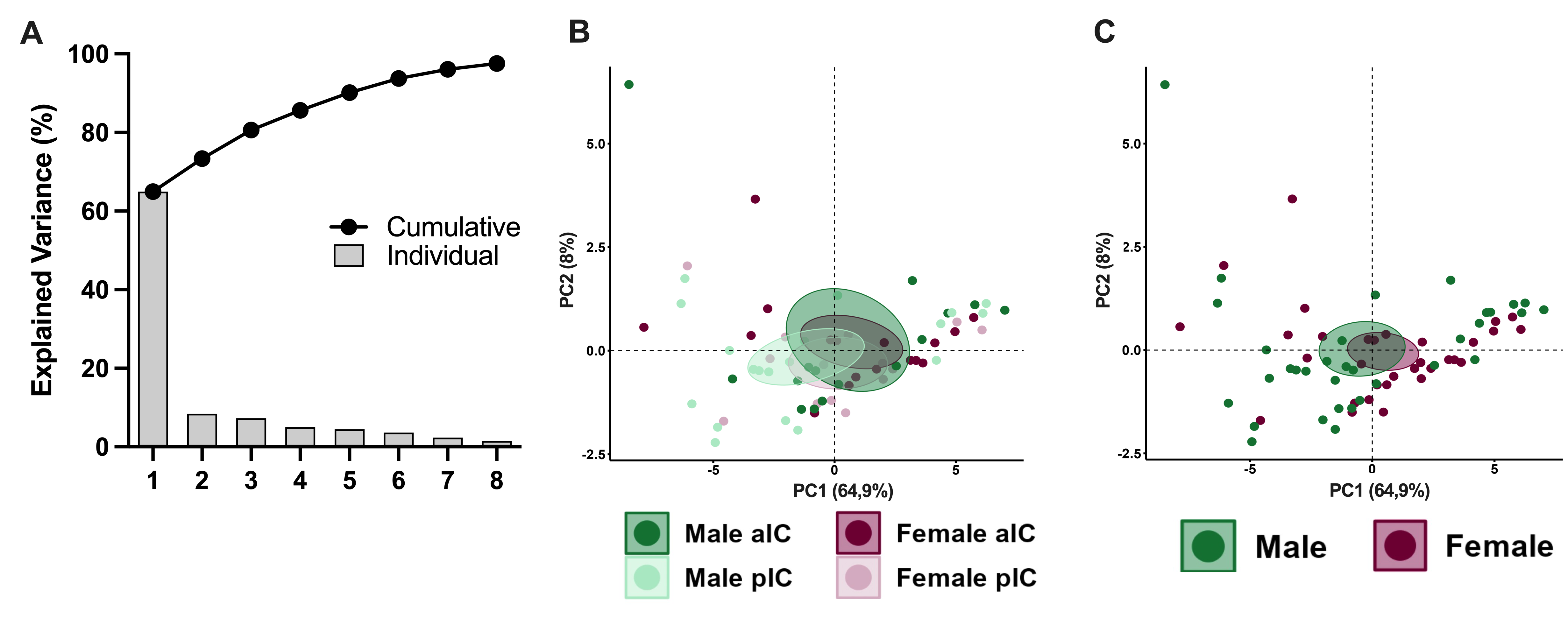

### Supplemental Figure 4 Iezzi

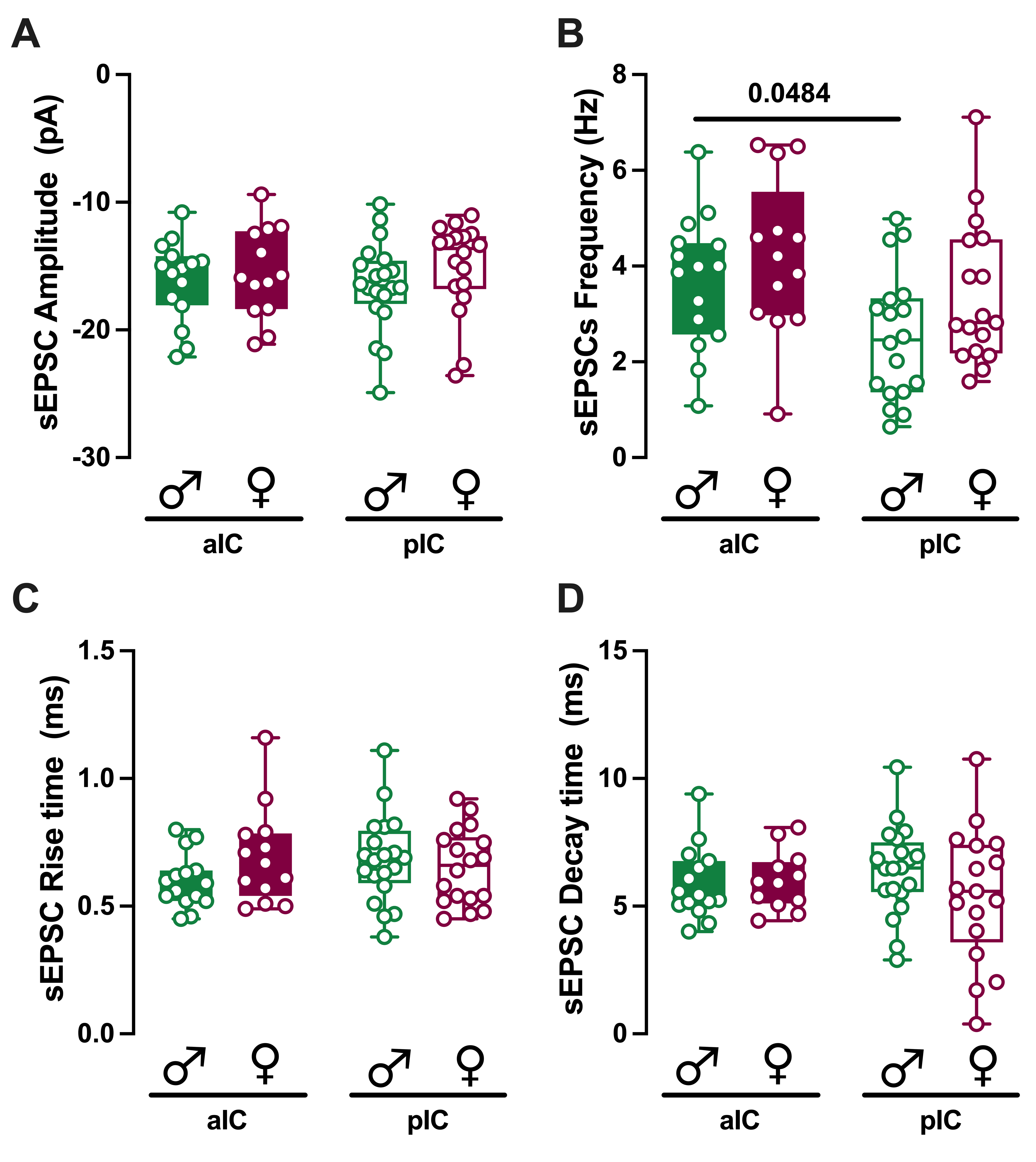

### Supplemental Figure 5 Iezzi

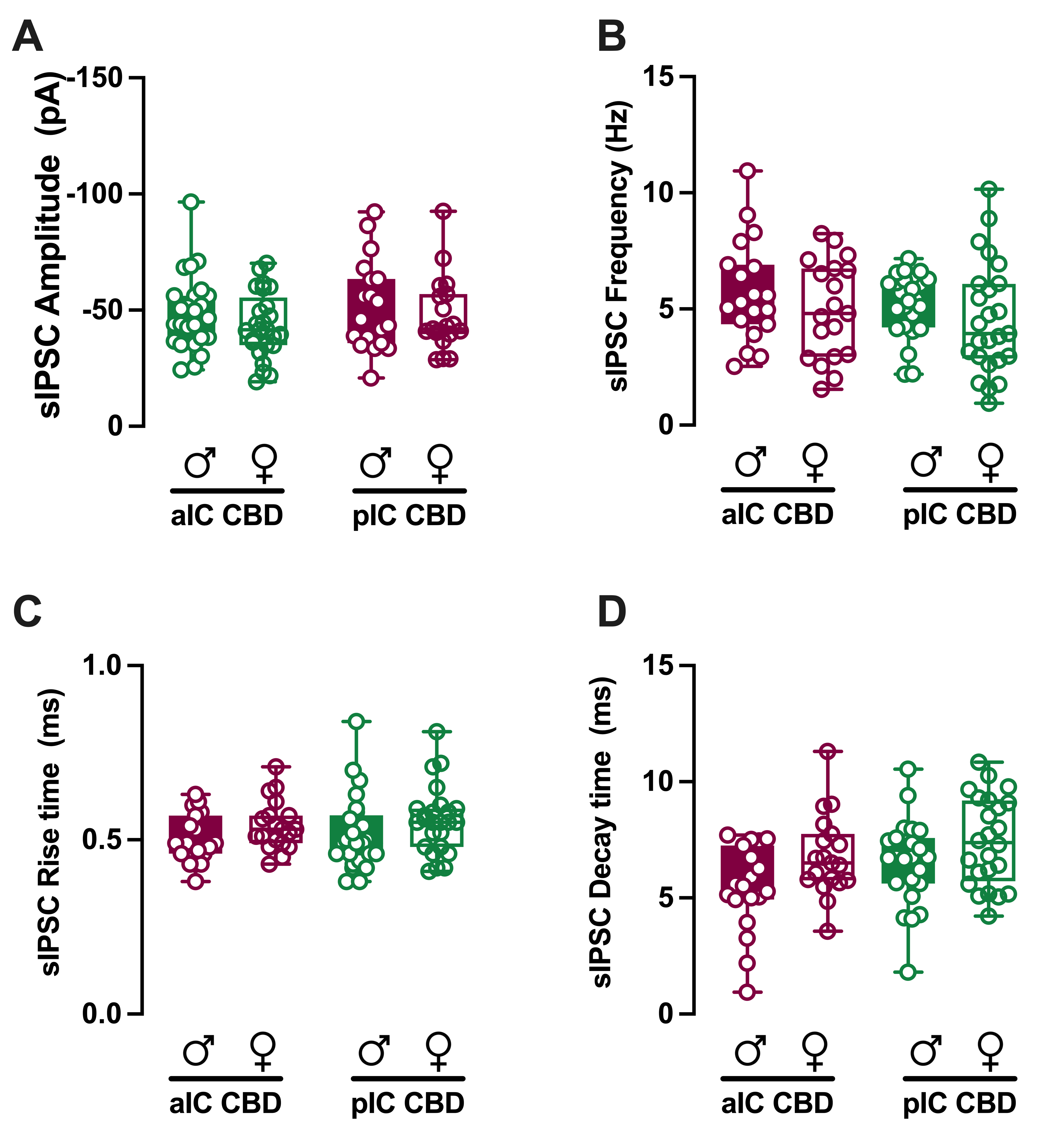
